# Selective connectivity limits functional binocularity in the retinogeniculate pathway of the mouse

**DOI:** 10.1101/2020.10.14.339747

**Authors:** Joel Bauer, Simon Weiler, Martin Fernholz, David Laubender, Volker Scheuss, Mark Hübener, Tobias Bonhoeffer, Tobias Rose

## Abstract

Eye-specific segregation of retinal ganglion cell (RGC) axons in the dorsal lateral geniculate nucleus (dLGN) is considered a hallmark of visual system development. However, a recent anatomical study showed that nearly half of the neurons in dLGN of adult mice still receive input from both retinae, but functional data about binocularity in mature dLGN is conflicting. Here, we found that a variable but small fraction of thalamocortical neurons is binocular *in vivo*. Using dual-channel optogenetics *in vitro* we correspondingly found that dLGN neurons are dominated by retinogeniculate input from one eye only, although most neurons also received small but detectable input from the non-dominant eye. Anatomical overlap between RGC axons and dLGN neuron dendrites did not explain this strong bias towards monocularity. Our data rather suggest that functional input selection and refinement, leaving the remaining non-dominant eye inputs in a juvenile-like state, underlies the prevalent monocularity of neurons in dLGN.

## Introduction

The retinogeniculate pathway of mice is a widely studied model for the segregation and refinement of neuronal connectivity in sensory systems^1^. However, recent studies have provided seemingly conflicting evidence about the degree of convergence and input selection at the level of single cells in mouse dorsal lateral geniculate nucleus (dLGN)^2–8^. Specifically, the degree to which neurons are still binocular in the mature dLGN is currently unclear^2,9–13^.

In the mammalian brain, retinal ganglion cells (RGCs) connect to neurons in the dLGN in a highly structured manner, and RGC afferents are separated by visuotopy, RGC type, and eye of origin^1^. In carnivorans and primates, thalamocortical (TC) neurons with distinct functional properties segregate into well-defined layers that receive retinotopically arranged input from different RGC types^14,15^. Binocular responses of individual TC neurons are rare in cats^16,17^ and are all but absent in the main magno- and parvocellular dLGN layers of primates^18^. With the exception of binocular neurons in koniocellular layers^19^, suppressive binocular modulation is the dominant thalamic interaction between the segregated eye-specific channels in primate dLGN^18,20^. TC neurons in these species are therefore generally thought to relay eye-specific visual information to the binocular visual cortex (bV1) via parallel and largely unmodified streams^14^.

The mature dLGN of rodents lacks clear lamination, but also here, eye-specific RGC projections are distinctly segregated^21–23^. Large-scale eye-specificity of RGC afferents in dLGN is the result of a developmental process which consists of a transition from diffuse retinogeniculate connectivity to an ordered network with clear eye-specific termination zones^23–25^. On the fine-scale synaptic level, electrophysiological experiments using randomly stimulated RGCs have shown that large-scale axonal segregation is accompanied by input selection and synaptic refinement, such that only a small number of RGCs contribute to the firing of a TC neuron in the mature dLGN^3,4,7,26^. The establishment of the mature eye-specific pattern is additionally reflected by the developmental increase in the contribution of AMPA receptors (AMPAR) over NMDA receptors (NMDAR) at the retinogeniculate synapse^3,27^.

Functional convergence of the two eyes onto dLGN neurons of mice is classically thought to be largely eliminated during the first postnatal weeks^12,28^. Consistent with this, calcium imaging studies of dLGN TC neuron afferents in V1 of adult mice found that the majority of TC boutons are exclusively driven by only one eye^10,11^. However, other recent studies have provided evidence that connections from both eyes to dLGN neurons may be much more wide-spread. Two electrophysiological *in vivo* studies showed that most dLGN neurons in the medio-dorsal part subserving the binocular field-of-view respond to stimulation of either eye^9,29^. Likewise, a report using mono-synaptic retrograde rabies tracing not only showed that many tens of RGCs can converge onto a single dLGN neuron, but also that close to half of all tested dLGN cells receive input from RGCs of both retinae^2^. Together with the finding that mouse dLGN neuron dendrites frequently cross the border between the eye-specific axonal projection zones^30^, these data suggest that the binocularity of a subset of geniculate neurons originates from direct retinal input rather than from cortical feedback projections. Therefore, there is an apparent discrepancy between structural tracing data, functional *in vitro* results, and several recent *in vivo* recording studies.

The present study was designed to resolve these inconsistencies and provide a better understanding of sensory integration and eye-specific input selection at the level of single cells in the mature mouse dLGN. We found that even though a large fraction of dLGN neurons received detectable functional input from RGCs of both retinae *in vitro*, dLGN neurons were largely dominated by input from one eye only. Morphometric analysis showed that indiscriminate dendritic sampling from segregated eye-specific RGC axons alone cannot explain this prominent functional monocular bias. Rather, highly specific input selection seems to be the main driver of monocularity. Importantly, we found that the non-dominant eye synapses had prominently reduced AMPAR / NMDAR ratios, reminiscent of a juvenile state. We conclude that fine-scale synaptic selection and refinement of eye-specific input strength is likely to be the driving factor behind the surprising degree of monocularity at the retinogeniculate synapse.

## Results

### A small fraction of thalamocortical neurons is binocular *in vivo*

Recent *in vivo* reports on TC neuron activity provided a wide range of estimates of functional binocularity in mouse dLGN^9–11,29^. These studies used different recording techniques (single unit electrophysiology vs. calcium imaging), different visual stimulus types (full-field luminance stimuli vs. full- and sub-field moving gratings), different analysis methods and responsiveness criteria, and different dLGN neuron sampling (across dLGN vs. recordings exclusively from TC afferents in L1 of bV1). To obtain a more consistent estimate of thalamic functional *in vivo* binocularity we therefore aimed to systematically explore the effect of all of these factors by recording from TC afferents in bV1 using two-photon calcium imaging (Fig. 1a,b).

**Fig. 1:**
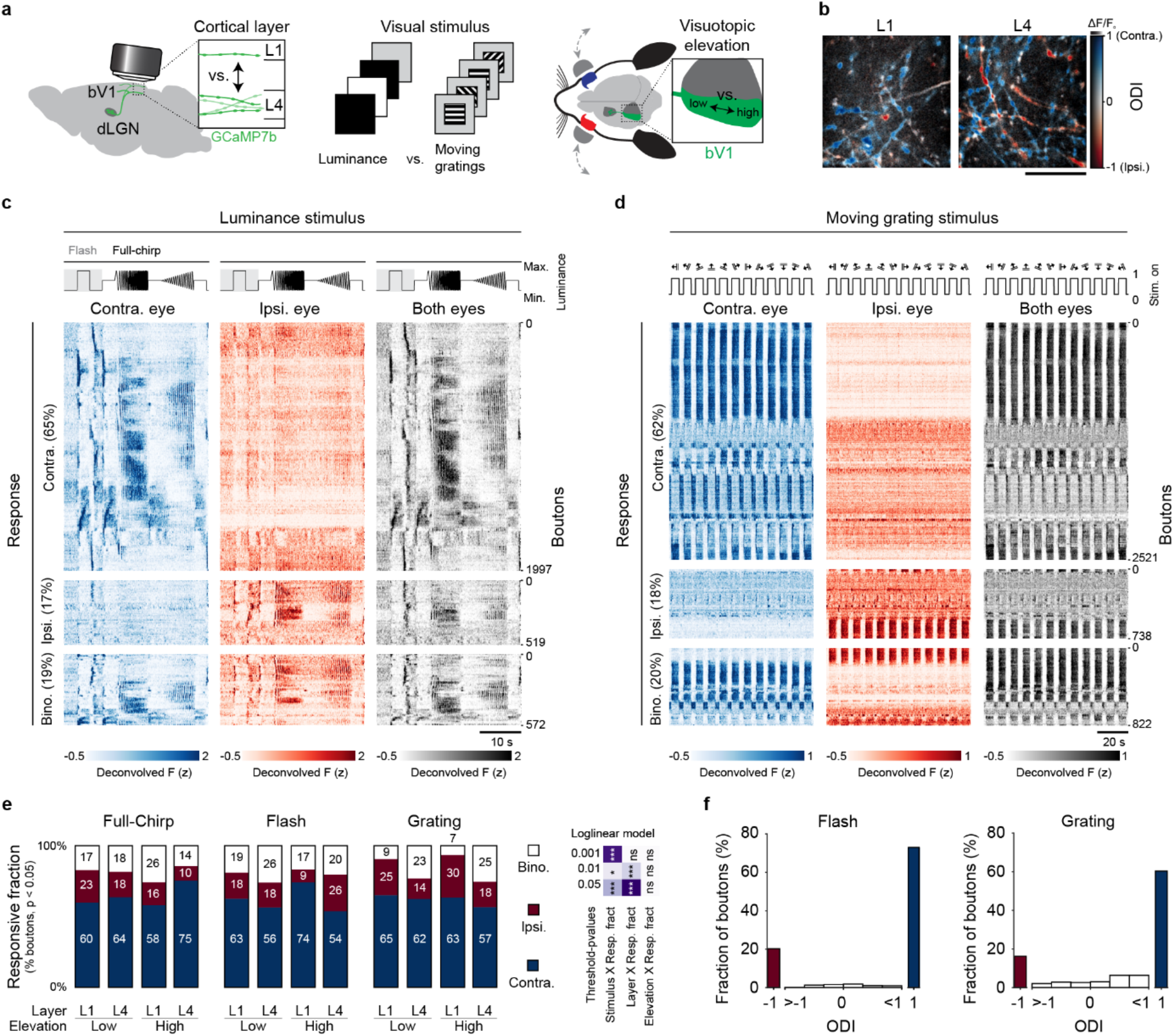
A variable but small fraction of thalamocortical neurons is binocular *in vivo*. **a,** Schematic of imaging approach across cortical thalamorecipient layers 1 (L1) and 4 (L4), using both full-field luminance and moving grating stimuli, recorded at two different visuotopic elevations (low = 0°, high = 20° elevation). **b,** Examples of color-coded response maps of individual dLGN boutons in L1 (left) and L4 (right). Ocular dominance is depicted as the pixel-wise peak fluorescence change in response to ipsi- and contralateral eye preferred grating presentation. Red hues indicate ipsilateral dominance (ODI < 0) and blue hues indicate contralateral dominance (ODI > 0). **c,** Activity of 3088 boutons (N = 4 mice; including significant positive and negative responses) across layers and elevations in response to contralateral, ipsilateral, and binocular full-field luminance stimulation, grouped by eye-specific responsiveness and sorted by similarity of the joint stimulus-aligned data (boutons with highly correlated responses are plotted next to each other). **d,** As in c (n = 4081 boutons, N = 4 mice), but in response to moving grating stimulation. **e,** Eye-specific response fractions across all conditions (including significant positive and negative responses). Right: p-value table of log-linear model of eye-specific response partitioning, comparing select pairwise interaction vs. non-interaction main effect model (residual deviance = 781, df = 29): Stimulus X Response fraction (residual deviance = 753, df = 25), *χ*^2^(2) = 27.9, p < 0.001; Layer X Response fraction (residual deviance = 724, df = 27), *χ*^2^(2) = 57.7, p < 0.001; Elevation X Response fraction (Residual deviance = 776, df = 27), *χ*^2^(2) = 4.9, p = 0.09; log likelihood-ratio model comparison using chi-squared test; see Supplementary Fig. 1c,d for data on different responsiveness criteria and further statistical analysis). **f,** Single-bouton ODI distributions based on full-field flash responses (left, mean |ODI| = 0.95 ± 0.18, positive responses only) and moving grating stimuli (right, mean |ODI| = 0.9 ± 0.23). Colored ODI histogram bins indicate class definitions for exclusively contralateral (blue), binocular (white) and exclusively ipsilateral (red) boutons. Scale bar in b: 30 μm.

We sparsely expressed the genetically-encoded calcium indicator (GECI) GCaMP7b in dLGN of *Scnn1a-Tg3-Cre* mice with custom adeno-associated viruses and compared responses to eye-specific moving gratings^10,11^ and eye-specific full-field luminance stimuli^7^. The latter consisted of an initial full-contrast flash^9,29^ followed by frequency- and contrast-modulated luminance changes^7^. We recorded the stimulus-evoked activity of individual TC neuron boutons in both superficial (L1, 10-90 μm below pia) and deep cortical layers (L4, 320-450 μm below pia), in response to stimuli presented in the central visual field and at higher elevation.

Fig. 1c,d shows the activity of TC boutons pooled across thalamocortical projection layers and elevations in response to contralateral, ipsilateral, and binocular full-field luminance (Fig. 1c) and moving grating stimulation (Fig. 1d). Using a lenient single criterion (p < 0.05) for neuronal responsiveness (see methods) we found that, pooled across all conditions and stimuli, 21% of TC neurons were significantly activated or suppressed by stimulation of either eye. This number reduced to 7% with more stringent selection criteria (p < 0.001; Supplementary Fig. 1d), underlining that absolute values of binocularity prominently depend on statistical selection criteria. Multivariate analysis (loglinear model, see methods) showed a significant interaction of stimulus type with eye-specific partitioning (Fig. 1e, Supplementary Fig. 1c) and pooled data showed a larger fraction of binocular neurons for the initial flash part of the full luminance stimulus than for other stimuli (Supplementary Fig. 1d). However, the maximum deviation of the binocular fraction across stimuli was small (2.6% across responsiveness criteria, conditions pooled, see methods) and inconsistent, i.e. changing sign, when assessed with more stringent responsiveness criteria (Supplementary Fig. 1d).

Eye-specific input fractions may vary between input-specific partitions and visuotopy in the dLGN, which could lead to inhomogeneous binocularity of TC afferents. Mouse dLGN can be subdivided into a shell region receiving input from RGC neurons of largely contralateral eye origin and a core region that encompasses the majority of the contra- and ipsilateral eye RGC axon termination zones^22^. Shell neurons predominantly project to L1 of V1 whereas core neurons project most densely to L4^31^. In line with that notion, we found that eye-specific partitioning was different across cortical layers (Fig. 1e, Supplementary Fig. 1c,d). L4 showed a slightly larger binocular fraction on average, but again, this effect only held for relaxed threshold criteria and was of small magnitude (maximum deviation across responsiveness criteria = 3.2%; Supplementary Fig. 1d). Furthermore, the visuotopic organization of dLGN may lead to different eye-specific partitioning across the mouse visual field. That said, we found no difference in responsive fractions when presenting visual stimuli in the central visual field at lower versus higher elevations (Fig. 1e, Supplementary Fig. 1c).

Similar to cats and primates^18^, responses to moving grating stimulation under binocular viewing conditions were smaller than the dominant eye response to monocular stimulation (Supplementary Fig. 1 e). Both visuotopy and cortical projection layer had an effect on the degree of binocular response modulation. Responses in L1 were more suppressed than in L4 when stimulated through both eyes and the same was true for higher vs. lower elevations.

To obtain a better picture of the relative eye-specific response amplitudes, we calculated the ocular dominance index (ODI), exclusively based on stimulus-evoked increases in activity, in response to both full-field flashes and moving gratings. ODI value distributions for both stimuli were highly monocular. In contrast to eye-specific partitioning assessed with lenient responsiveness criteria (Fig. 1e), full-field flash stimuli evoked slightly less binocular activity than sub-field moving grating stimuli (mean flash |ODI| = 0.95 ± 0.18 vs. mean grating |ODI| = 0.9 ± 0.23; p = 3e-14; Mann-Whitney U test; Fig. 1f, Supplementary Fig. 1f,g), further underlining the complexity of binocular interactions in mouse dLGN.

In summary, the degree of eye–specific partitioning is affected by all tested conditions except retinotopic elevation. However, the effect sizes are small and often inconsistent when assessed with different responsiveness criteria. Importantly, TC neuron binocularity is consistently low.

### A dual-channel input mapping approach for studying crosstalk-free eye-specific retinogeniculate convergence

The low level of functional binocular *in vivo* responsiveness we and others^11^ observe is seemingly at odds with a recent neuronal tracing study showing a much larger fraction of dLGN neurons receiving anatomical input from both retinae^2^.

To understand how apparent anatomical convergence at the level of the retinogeniculate synapse results in relatively moderate functional binocularity, we implemented a dual-channel optogenetic *in vitro* approach to measure functional eye-specific input to individual dLGN neurons in adult mouse brain slices. We virally expressed two channelrhodopsin variants in RGCs of either eye using ChrimsonR and Chronos, which have minimal spectral overlap, to stimulate retinogeniculate axons with eye-specificity (Fig. 2a)^32^. The tagging of ChrimsonR with tdTomato (tdT) and of Chronos with EGFP allowed visualization of RGC afferent projection zones within dLGN^21^ during the electrophysiological recordings and post-hoc in cleared brain slices using confocal microscopy (Fig. 2a).

**Fig. 2:**
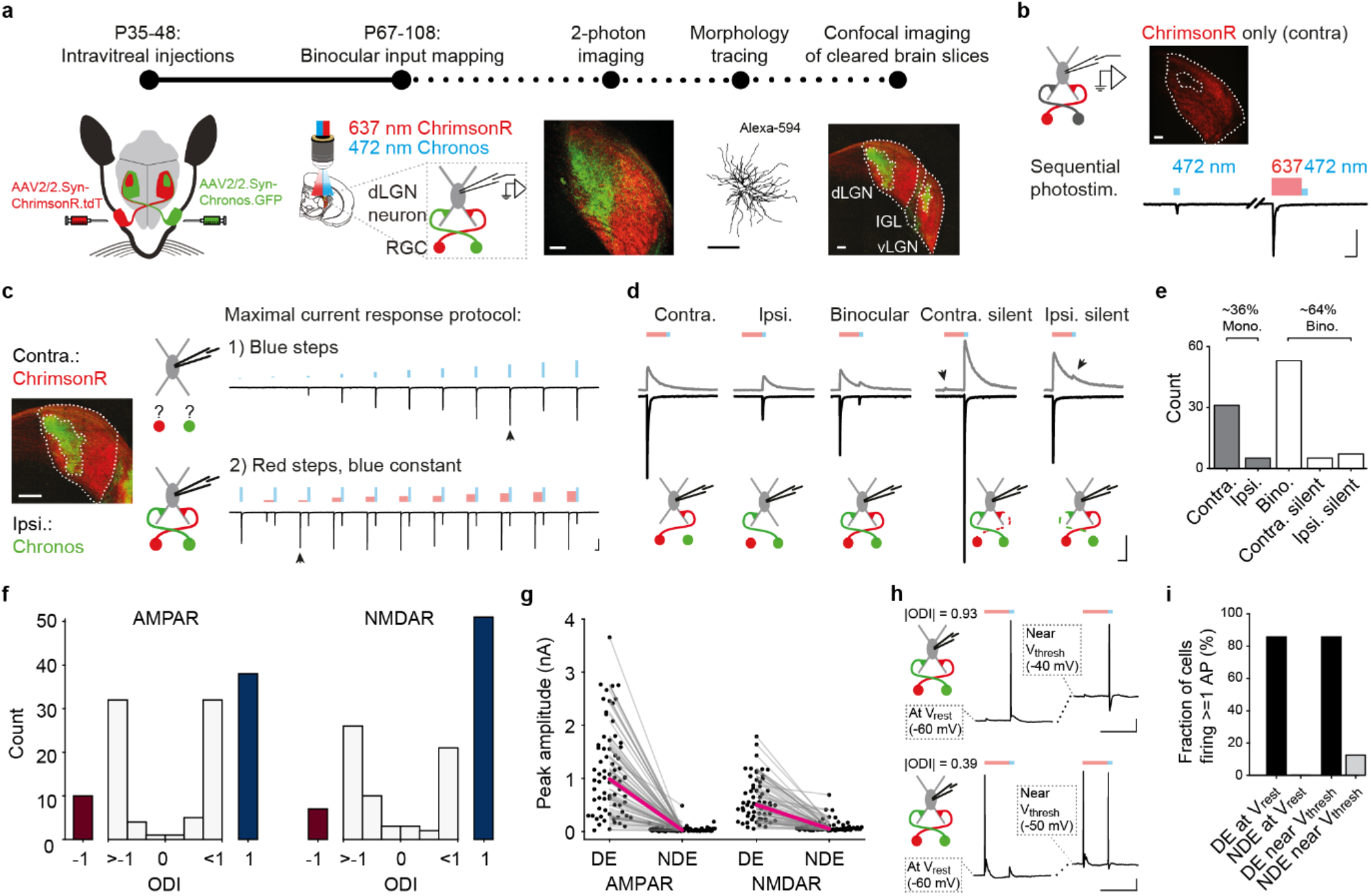
Eye-specific functional input mapping reveals prominent retinogeniculate convergence but limited functional binocularity. **a,** Experimental pipeline (from left to right): Red- and blue-light excitable ChrimsonR/Chronos (are expressed in RGCs). Individual dLGN neurons are recorded and ChrimsonR+/Chronos+ retinogeniculate axons are optogenetically stimulated by red and blue light using sequential photostimulation. Maximum intensity projection of expression pattern of ChrimsonR-tdT and Chronos-EGFP in RGC axonal terminals within the dLGN obtained by two-photon imaging. Reconstructed morphology of an Alexa-594-filled dLGN neuron. Maximum intensity projection of cleared brain slice using confocal imaging. **b,** Suppression of crosstalk using sequential photostimulation: maximum intensity projection of expression pattern of ChrimsonR-tdT in RGC axonal terminals within dLGN (top). PSC (from dLGN neuron clamped to −70 mV) evoked by stimulation at 473 nm (50 ms, left trace; irradiance: 5.1 mW/mm^2^) and by sequential photostimulation with stimulation at 637 nm (250 ms) and 473 nm (50 ms, right trace, irradiance 473 nm: 5.1 mW/mrtf; irradiance 637 nm: 3.4 mW/mrtf). **c,** Maximum intensity projection of RGC axonal terminals expressing ChrimsonR and Chronos within dLGN. Maximal current response protocol: 1) evoked currents of recorded dLGN neuron upon stimulation at 473 nm using 11 steps with increasing irradiance (pulse duration: 50 ms, ISI: 10 s). 2) Evoked currents of the same dLGN neuron using sequential stimulation at 637 nm (250 ms) and 473 nm (50 ms, ISI: 10 s). The red irradiance is increased across 11 steps while the blue irradiance is the irradiance that evoked the maximal response in protocol 1) (indicated with arrowhead in 2). Arrowhead in 2) indicates successful sequential photostimulation. **d,** Representative examples of observed eye-specific evoked PSC patterns at −70 (black) and +40 mV (gray). Silent inputs are only detectable at +40 mV (indicated with arrowheads). **e,** Quantification of main input categories (in **d**; n = 101 cells). **f,** AMPAR- (left) and NMDAR-based (right) ODI distribution within the adult dLGN (n = 123 cells). **g,** Evoked peak currents of the dominant (DE) and non-dominant eye (NDE) for AMPAR- (left, n = 75 cells) and NMDAR- (right) mediated responses for binocular cells (n = 65 cells). Medians indicated in magenta. **h,** Membrane voltage deflections upon eye-specific stimulation in two example binocular dLGN neurons. In both neurons the DE RGC stimulation evokes an action potential, whereas the NDE fails to trigger action potentials at resting (left). Depolarization close to action potential threshold (using current injection; right), in combination with NDE RGC stimulation, only triggers an action potential in the more binocular dLGN neuron displayed at the bottom. **i,** Quantification of **h** (n = 8 cells receiving input from both eyes). All image and morphology scale bars: 100 μm. Electrophysiology trace scale bars in **b-d**: 100 ms, 250 pA; and in **h**: 250 ms, 10 mV.

To quantitatively assess the maximum eye-specific input strength we aimed to activate the two axonal populations independently at saturating stimulation levels without spectral crosstalk. To this end, we adopted and modified a previously introduced sequential photostimulation approach^33^. This approach is based on the principle that prolonged stimulation with red light renders axons expressing the red-shifted opsin (ChrimsonR) refractory to a subsequent brief stimulation with blue light, which otherwise would elicit a non-specific response. Axons expressing the blue-shifted opsin, in turn, exclusively respond to the blue light pulse since they are essentially unresponsive to red light stimulation^32^.

First, we tested the feasibility of this approach in brain slices with RGC axonal projections expressing only ChrimsonR (Fig. 2b). We used whole-cell patch-clamp to record optically evoked retinogeniculate postsynaptic currents (PSCs) at −70 mV in individual dLGN neurons and blocked di-synaptic feedforward inhibition using bicuculline. As expected, light stimulation at 473 nm evoked large amplitude PSCs, similar to previous channelrhodopsin-assisted circuit mapping (CRACM) experiments, in dLGN^4^ (Fig. 2b left trace, and Supplementary Fig. 2a). However, photostimulation of retinogeniculate axons expressing ChrimsonR with 637 nm light for 250 ms, immediately followed by illumination at 473 nm for 50 ms, did not evoke a second PSC in the same dLGN neurons, confirming the viability of the sequential photostimulation approach in our preparation (Fig. 2b, bottom). We chose a 250 ms long red light stimulus to obtain efficient crosstalk suppression and to temporally separate the responses evoked by red and blue light stimulation (Supplementary Fig. 2b,c). At this pulse duration we observed prominent rundown of the peak PSC amplitude (Supplementary Fig. 2d,e). However, rundown was completely prevented when pairing the red pulse with blue light stimulation (Supplementary Fig. 2d,e).

Therefore, this approach provides a novel procedure to conduct crosstalk-free mapping of eye-specific RGC input to dLGN neurons in the adult mouse.

### Prominent eye-specific retinogeniculate convergence but limited functional binocularity

The sequential photostimulation approach allowed us to study functional retinogeniculate ocular convergence in the dLGN and to precisely quantify relative eye-specific input strength, including the contributions of AMPARs and NMDARs to these responses. To achieve this, we used two stimulation protocols with increasing laser intensity for blue and red pulses in the same cell (Fig. 2c, see Methods). We recorded neurons across the entire dLGN with a focus on the ipsi- and the surrounding contralateral projection zone. Example traces of AMPAR- and NMDAR-mediated inputs are shown in Fig. 2d.

We observed two main input categories of dLGN neurons: Purely monocular (i.e. contra- or ipsilateral) neurons and neurons receiving input from both eyes. This included both ipsi- and contralaterally AMPAR-silent dLGN neurons (Fig. 2d) and a small fraction of neurons lacking NMDARs at retinogeniculate synapses (16%). Strikingly, 64% of the recorded dLGN neurons were binocular when considering NMDAR inputs (Fig. 2e). To quantify the degree of ocular dominance, we computed the AMPAR- and NMDAR-based ODI using the peak amplitudes evoked by stimulation of the inputs from either eye (see methods). The ODI distributions for AMPAR and NMDAR responses revealed that although there were numerous binocular cells, most of them were strongly dominated by input from one eye (AMPAR mean |ODI| = 0.91 ± 0.014). Only a minority of cells had an ODI close to 0 (Fig. 2f). Inspection of binocular cells showed that the AMPAR-mediated inputs of the dominant eye were on average 36× larger compared to the non-dominant eye (Fig. 2g, left, Supplementary Fig. 3b). The same was true for NMDAR-mediated inputs, albeit less pronounced (Fig. 2g, right, Supplementary Fig. 3c). Both the contra- and ipsilateral eye could provide dominant input (71 vs. 48 cells) and eye-dominance was identical when assessed by AMPAR and NMDAR responses for 122 out of 123 cells (Supplementary Fig. 3d).

Finally, we probed whether the inputs from the dominant and non-dominant eye could independently trigger action potentials in the same binocular dLGN neurons. For this, we measured evoked postsynaptic membrane voltage deflections upon sequential photostimulation in binocular dLGN neurons (Fig. 2h). Whereas the input from the dominant eye evoked action potentials in all but one dLGN neuron, non-dominant eye stimulation failed to trigger action potentials at resting membrane potential (Fig. 2i, left). Even when injecting current to depolarize the cell only 1 out of 8 binocular neurons fired action potentials upon non-dominant eye stimulation (Fig. 2i, right).

Taken together, our data shows that even though most neurons received detectable input from both eyes, dLGN neurons are by and large functionally dominated by input from one eye only.

### Mixed ocular dominance across dLGN regions and visuotopic space

Our *in vivo* and *in vitro* data showed that on average the degree of functional binocularity was relatively low in dLGN. However, eye-specific RGC input convergence could vary across dLGN. To elucidate the 3D distribution of eye-dominance across dLGN, we registered the cells in our dataset to the Allen Common Coordinate Framework (ACCF; Fig. 3a and Supplementary Video 1). As expected, we found that contra- and ipsilaterally dominated dLGN neurons were more prevalent in their respective eye-specific projection zones^21^. Nevertheless, based on their dominant input source, 24% of cells were located outside their expected projection zones (Fig. 3b,c), indicating that contra- and ipsilaterally dominated dLGN neurons were only partially segregated. However, the ACCF projection zone boundaries represent an average across animals. Therefore, deviations between individuals, as well as imperfect segregation between the projection zones, may partially account for some cells having opposite eye-dominance to the ACCF region their somas were located in.

**Fig. 3:**
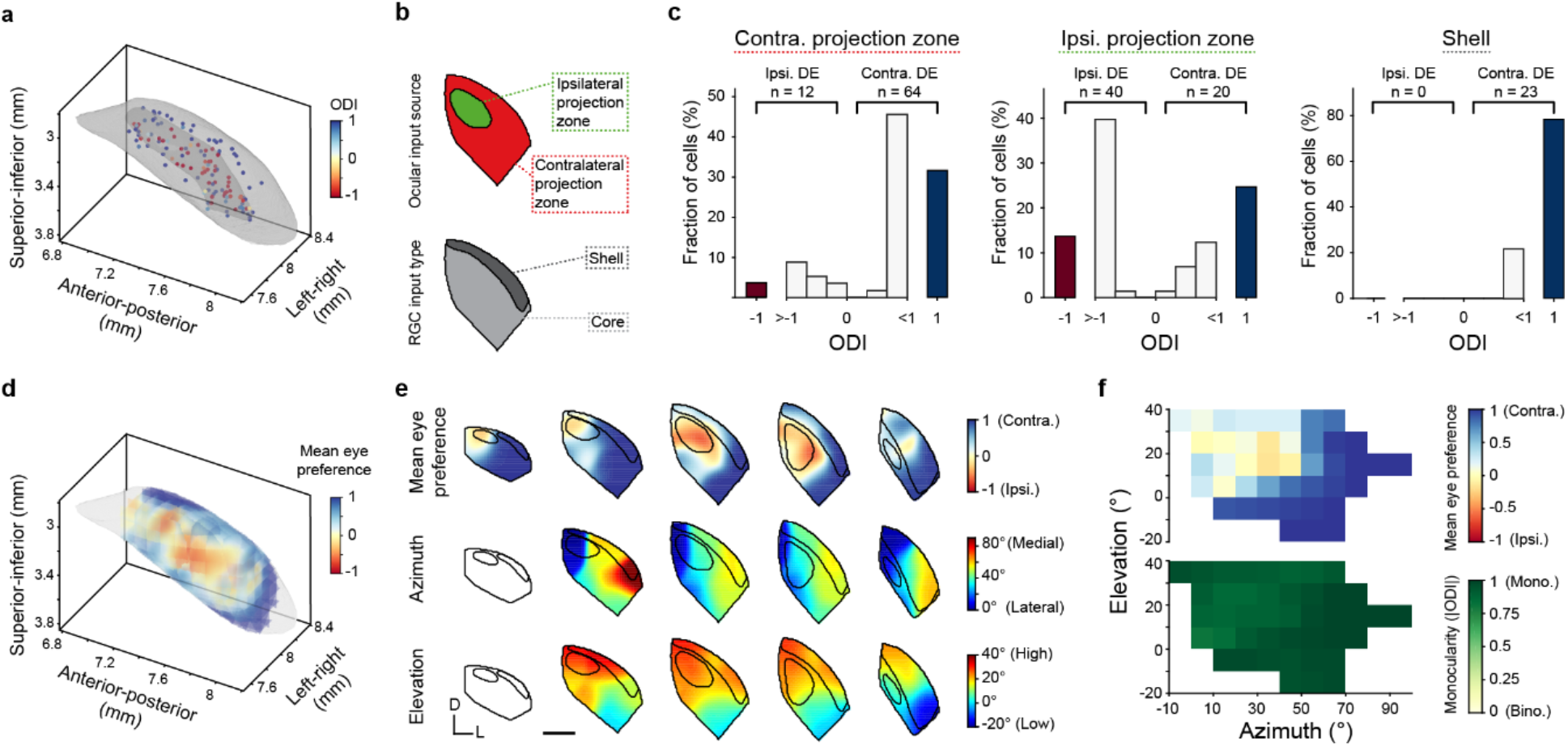
Ocular dominance across dLGN regions and visuotopic space. **a,** Position of recorded dLGN neurons within the ACCF, color coded by AMPAR-based ODI. dLGN and ipsilateral projection zone borders are indicated in gray (n = 136 cells). **b,** Anatomical subdivisions of dLGN. **c** ODI histograms of dLGN neurons within the contra- (n = 76 cells) and ipsilateral (n = 60 cells) projection zones of the core, and shell region (n = 23 cells), as defined by the ACCF. Red: exclusively responsive to ipsilateral input, blue: exclusively responsive to contralateral input, white: cells receiving input from both eyes. The contralateral projection zone of the dLGN core contains more contralaterally dominated cells compared to the ipsilateral projection zone (n = 136 cells, Cross-tab test *χ*^2^(1) = 36.8, p < 0.001). We found no ipsilaterally dominated cells in the shell region (n = 23 cells). **d,** Mean eye dominance map. ODI values in **a** are converted to eye dominance (−1 for ipsilaterally dominated, ODI < 0; 1 for contralaterally dominated, ODI > 0). Interpolation was performed using a 3D Gaussian kernel with 75 μm standard deviation. dLGN border indicated in gray. **e,** Multiple coronal sections, along the rostro-caudal axis, through 3D interpolated map of mean eye dominance in **d** and ACCF aligned visuotopy (data generously provided by the Niell lab, Piscopo et al.^34^). dLGN, ipsilateral projection zone and shell region outlines taken from ACCF. **f,** Mean eye dominance and mean monocularity (|ODI|) across visual space based on data in **e**. Scale bar in **e**: 200 μm.

When comparing cells in the shell and core regions of dLGN we found that cells recorded in the latter showed mixed eye-specific selectivity. Neurons in the dLGN shell, however, were exclusively contralaterally dominated, although some received small amounts of ipsilateral input (Fig. 3b,c). This is likely explained by the dendrites of neurons in the shell having little to no access to ipsilateral inputs^22^.

In the dLGN, RGC afferents are not only separated by eye of origin, but also by visuotopy. To examine the relationship between ocular dominance and visuotopy, we first generated a 3D map of average eye preference (Fig. 3d and Supplementary Video 1) and compared this to similarly re-analyzed *in vivo* dLGN visuotopy data, generously provided by the Niell lab^34^ (see methods; Fig. 3e). From these 3D maps, we then derived the average eye-dominance in relation to visuotopy (Fig. 3f). We found that the dLGN region representing the upper central visual field processed by both retinae^35^, contained a mixed population of contra- and ipsilaterally dominated neurons. Furthermore, by mapping absolute ODI to visuotopy we found that the level of monocularity was high throughout the visual field (Fig. 3f).

We therefore conclude that monocularity dominates throughout all regions of the dLGN and that the binocular visual space is represented by a mixed population of largely monocular neurons, similar to our *in vivo* data (Fig. 1).

### Three potential mechanisms leading to functional monocularity

Given the surprisingly low level of eye-specific RGC input convergence across dLGN, we next asked which mechanisms could underlie this wide-spread bias. Such functional monocularity at the retinogeniculate synapse could arise by three putative mechanisms: 1) Axon segregation: The anatomical segregation of RGC axons in the dLGN in combination with the average dendritic sampling area of dLGN neurons may lead to most cells accessing inputs from only one eye (Fig. 4a). 2) Dendritic orientation: dLGN neuron dendrites may be oriented with respect to local RGC innervation, thereby biasing the sampling of contra- or ipsilateral RGC axons (Fig. 4b). 3) Input selection and synaptic refinement: dLGN neurons may preferentially form synapses with axons from one eye even if inputs from both are available and adjust their strength in an eye-specific fashion (Fig. 4c).

**Fig. 4:**
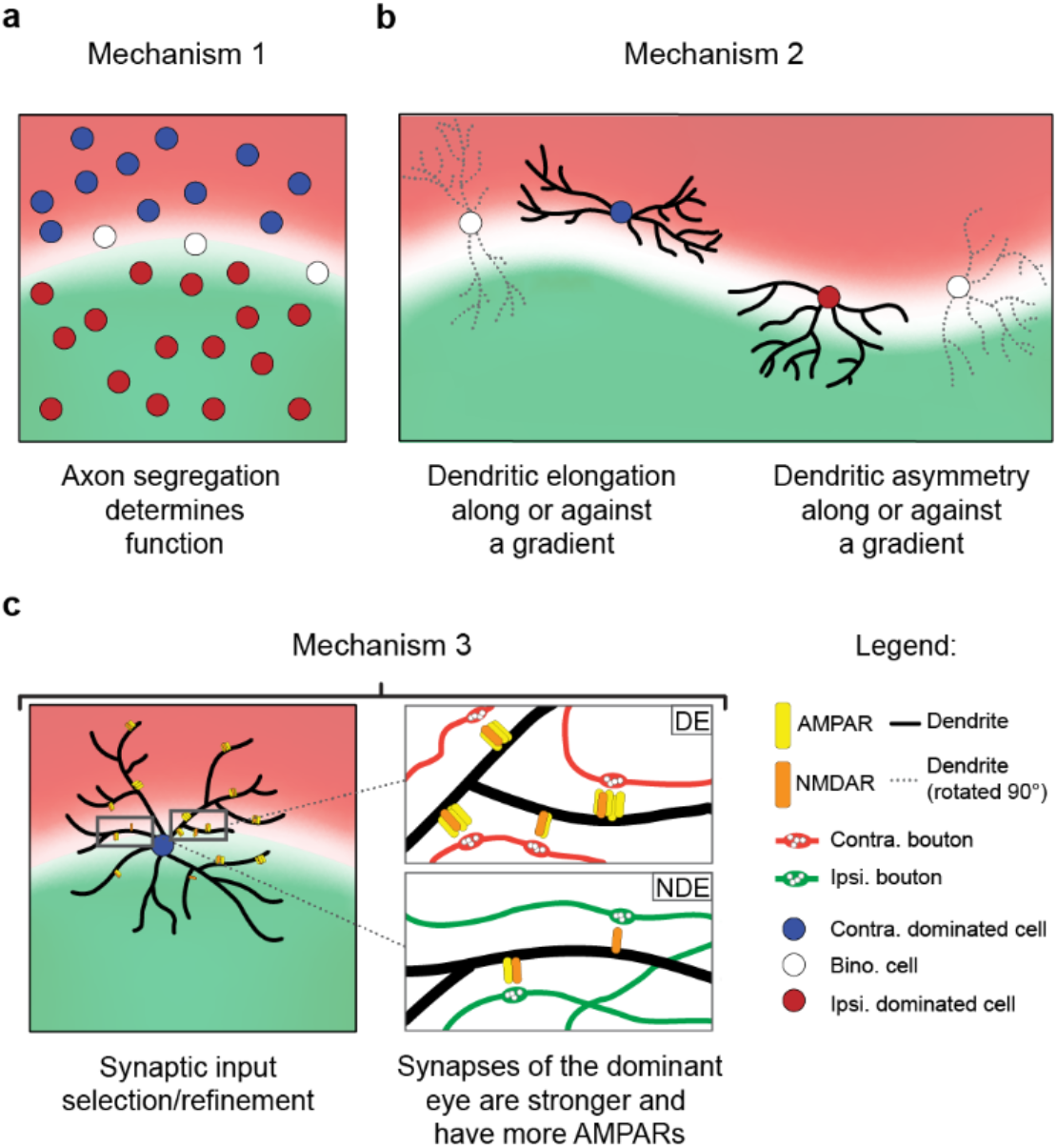
Three possible mechanisms leading to a dLGN neuron population bias towards monocularity. Three putative mechanisms underlying the observed monocular bias: **a,** Axon segregation: as most cells lie within either the contra- or the ipsilateral projection zones dLGN neurons will primarily have access to inputs from only one eye. **b,** Dendritic organization: the organization of dLGN neuron dendrites leads to biased sampling of available axons. **c,** Synaptic selection and refinement: largely irrespective of input availability, dLGN neurons preferentially form more numerous and stronger synapses with only one of the axon pools.

### Axon segregation does not explain functional monocularity

Depending on the geometry of the projection zones as well as the input sampling area of dLGN neurons, there could be an inherent bias towards neurons being dominated by input from only one eye (Fig. 4a). If this were the case, then in principle there should be few locations in dLGN from which neurons have roughly equal access to both contra- and ipsilateral inputs, compared to locations from which they predominantly have access to only contralateral or only ipsilateral inputs. To test this, we determined the average radial histogram of the dendrite densities of Alexa-594 filled cells to construct a three-dimensional sampling mask (Fig. 5a,b). From the confocal image stacks of each cleared brain slice we calculated the normalized fluorescence difference between the red (contralateral) and green (ipsilateral) channels and binarized them to simulate conditions of perfect axonal segregation. We then measured the normalized difference between the contra- and ipsilateral projection zones accessible within the three-dimensional mask, to derive the radial mask based fluorescence difference (rFD). This was repeated at arbitrary locations in the cleared dLGN slices (Fig. 5c-e and Supplementary Fig. 4a). Given that the ipsilateral projection zone is much smaller than the contralateral zone, and is accordingly under-sampled, we only compared the number of sampling positions across all dLGN slices that are predominantly ipsilateral (rFD < −0.33), to those that are binocular (|rFD| <= 0.33). We found that the proportion between these two groups was roughly equal (normalized difference = 0.1, corresponding to an ipsilateral:binocular ratio of 1:0.82). In contrast, our estimate of the normalized difference of functionally ipsilateral (ODI < −0.33) and binocular (|ODI| <= 0.33) neurons across dLGN was strongly skewed in favor of ipsilateral neurons (normalized difference = 0.9, corresponding to an ipsilateral:binocular ratio of 1:0.05; Fig. 5f and Supplementary Fig. 4b-m). Hence, even under optimized conditions, axon segregation alone cannot explain the observed level of functional monocularity, if dLGN neuron morphologies are assumed to be radially symmetrical.

**Fig. 5:**
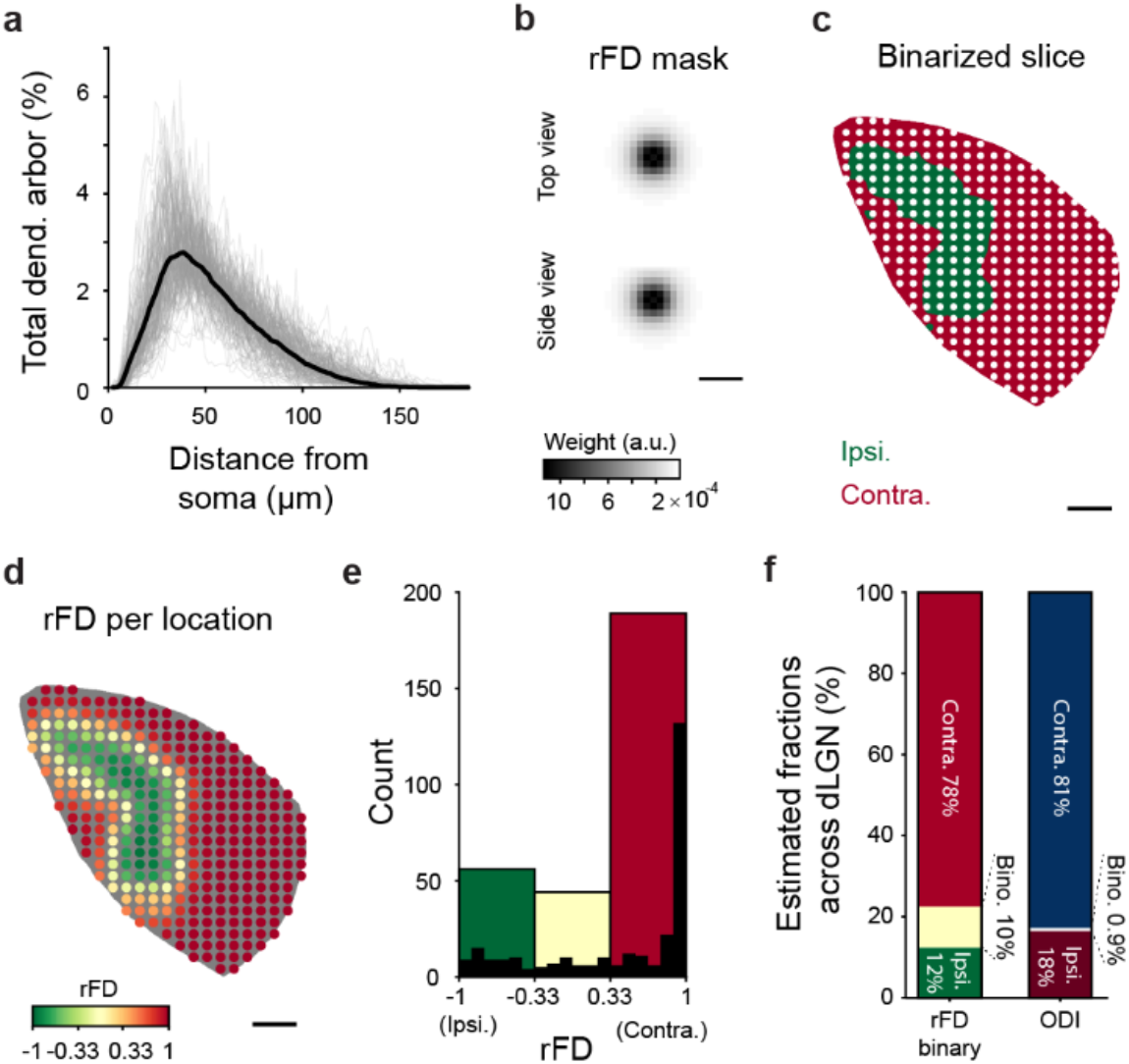
Axon segregation does not explain the monocular bias of dLGN neurons. **a,** Line histogram of dendrite density as a function of Euclidean distance from soma (gray: individual cells, black: mean of all cells). On average, 75% of the total dendritic length is located within a distance of 69 μm from the soma. **b,** 3D radial mask generated from mean distribution in **a. c**, Single plane of example dLGN slice after smoothing and binarizing confocal stack. White points: 40 μm spaced sampling positions. **d,** radial mask based normalized fluorescence difference (rFD) at each sampling positions, calculated using radial mask. **e,** rFD distribution of sampling positions in **d. f,** Estimated fraction of putative ipsilateral (rFD < −0.333; 12%), binocular (rFD >= −0.333 & <= 0.333; 10%) and contralateral (rFD > 0.333; 78%) sampling positions across dLGN (normalized mean across 21 slices), and estimated overall fraction of functionally ipsilateral (ODI < −0.333; 18%), binocular (ODI >= −0.333 & <= 0.333; 0.9%) and contralateral (ODI > 0.333; 81%) across dLGN (normalized across slices; n = 87 cells, 21 slices). All scale bars: 100 μm.

### Dendritic orientation does not account for functional monocularity

The dendritic morphologies of dLGN neurons are not always radially symmetrical, but rather show considerable variability^30^. This variability of morphologies could, in combination with RGC axon segregation, lead to the observed bias towards monocularity. We found that morphologies vary mainly along three uncorrelated measures: dendritic reach, elongation magnitude, and asymmetry magnitude (Fig. 6a and Supplementary Fig. 5a-c; see methods), although there was no evidence for discrete morphological groups (Supplementary Fig. 5d-e; similar to^5^ but see^30^). As the mean radial distribution of dendrites does not predispose to high monocularity (Fig. 5), dendritic reach was unlikely to have an effect. However, elongation, but not asymmetry, was correlated with and oriented orthogonally to the gradient of the eye-specific RGC axon innervation within 150 μm around each cell (Fig. 6a-c and Supplementary Fig. 5f). Hence, dendritic orientation has the potential to cause increased monocularity at the level of the dLGN neuron population (Fig. 4b).

**Fig. 6:**
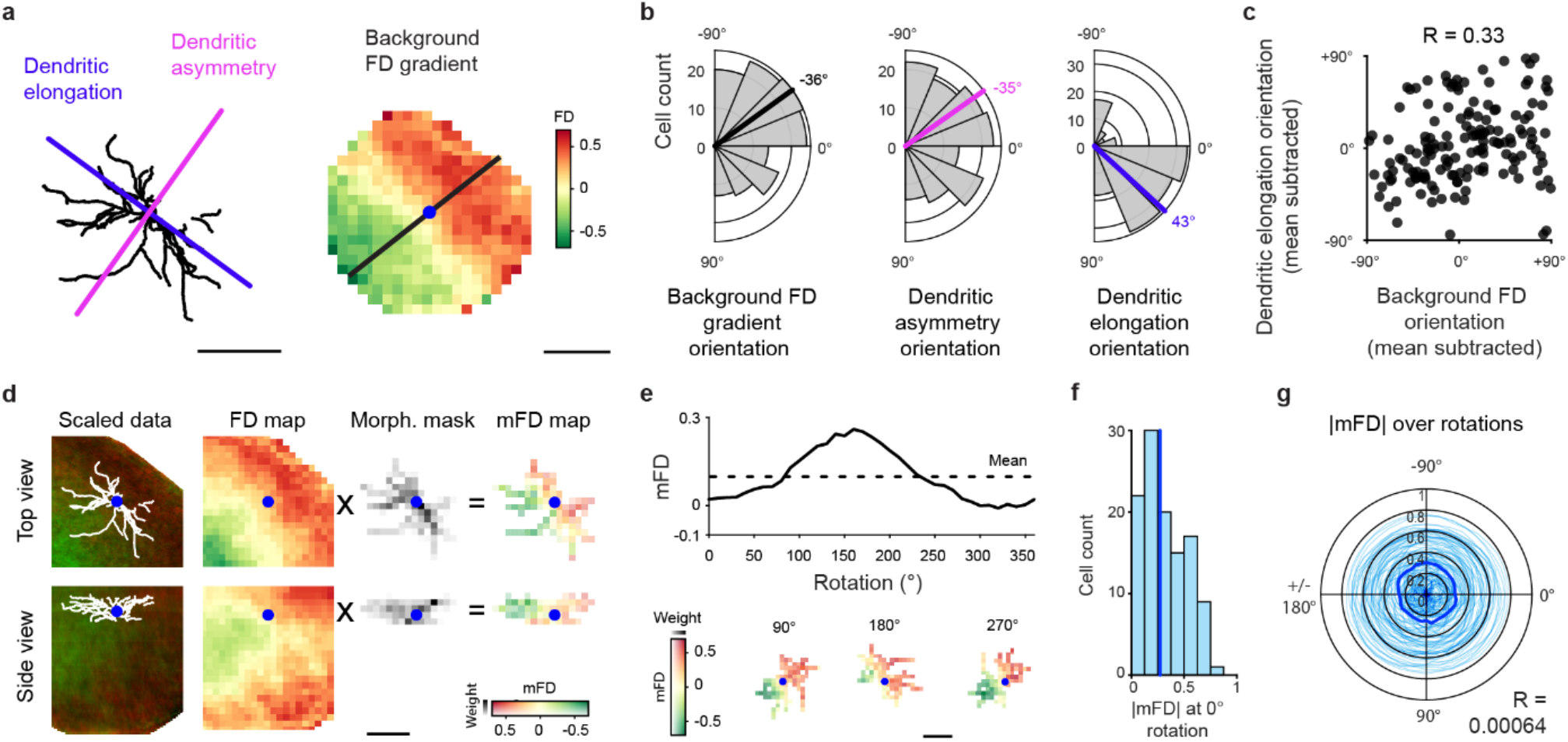
Dendritic orientation does not explain the monocular bias of dLGN neurons. **a,** Left: example cell morphology with orientation of asymmetry (magenta line) and elongation (blue line). right: z-projection of background normalized fluorescence difference (FD) within a 150 μm sphere around the cell. Black line: orientation of background FD gradient. Cell soma location indicated by blue circle. **b,** Distribution of background FD gradient, dendritic asymmetry and elongation orientation of dLGN neurons. Circular means indicated in black (−36°), magenta (−35°) and blue (43°) lines, respectively. **c,** Scatter plot of mean subtracted background FD gradient orientation vs. mean-subtracted elongation orientation (n = 152 cells, circular-circular Pearson’s correlation R = 0.33, p < 0.001). **d,** Process of calculating mFD for an example cell. Top row displays z-projection (top view), bottom row display x-projection (side view). Left to right: confocal stack with aligned morphology trace, FD pixel map, morphology mask, and morphology-based FD (mFD) pixel map. Cell soma location indicated by blue circle. **e,** Top: mFD of cell in **d** as it is rotated through 360° (solid line). Dashed line indicates the mean mFD across all rotations. Bottom: mFD pixel map from **d** at three rotations. **f,** Distribution of absolute mFD when morphologies are not rotated. Solid blue line indicates median (0.27). **g,** Absolute mFD of all cells when morphologies are rotated (light blue lines: single cells, solid blue line: median). There is no correlation between morphology rotation and absolute mFD (n = 114 cells, circular-linear Spearman’s correlation R = 0.00064, p = 0.28). All scale bars: 100 μm.

To investigate if the observed orientation of morphologies manifests into a meaningful bias towards monocularity, we estimated the relative difference in RGC axons available to each cell based on their individual morphology (morphology based fluorescence difference, mFD, Fig. 6d). As dendritic morphologies were rotated from their original alignment, the mFD of some cells fluctuated (Fig. 6e). If the orientation of elongation affects the overall level of monocularity in dLGN, then the absolute mFD of the population should be higher at the original orientation of morphologies and lowest when rotated orthogonally, resulting in a correlation between rotation and absolute mFD. However, we found no such correlation (Fig. 6f,g), suggesting that on the population level the actual orientation of morphologies did not lead to a more monocular mFD distribution. Morphological orientation may nevertheless play a role in other visual processes (e.g. compensation for visuotopic anisotropies; Supplementary Fig. 5g-i). We conclude that eye-specific RGC axon segregation in combination with dendritic organization of dLGN neurons has no effect on the level of monocularity.

### Synaptic selection and refinement results in functional monocularity

As both axonal segregation and biased sampling of axons by dLGN neuron dendrites were insufficient to explain functional monocularity, we determined synaptic sub-selection of eye-specific inputs by each dLGN neuron to be the most likely mechanism (Fig. 4c). Indeed, for most mFD values, we found dLGN neurons with either eye-dominance (Fig. 7a), even though availability of eye-specific axons to dendrites was a predictor of eye-dominance (Supplementary Fig. 6). Strikingly, neighboring dLGN neurons with near-identical mFD values and partially overlapping dendrites within the same slice sometimes showed opposite eye-dominance (Fig. 7b). Nevertheless, there was a roughly linear relationship between mFD and the probability of contra- or ipsilateral input dominance (Fig. 7c). Taken together, dLGN neurons selectively form stronger and / or more numerous synapses with one of the two input sources, selected in a probabilistic manner dependent on the relative availability of both inputs.

**Fig. 7:**
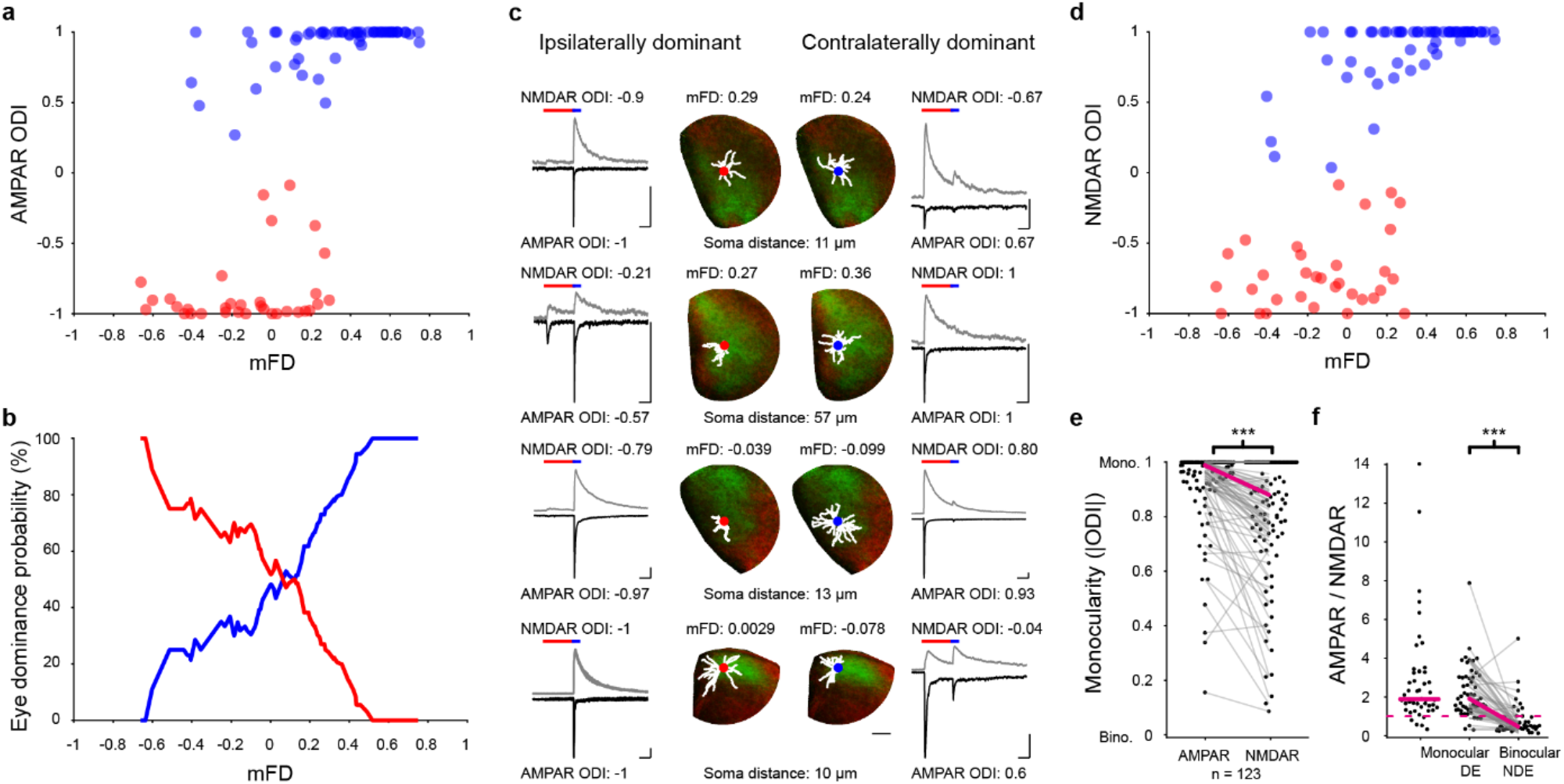
Synapse selection and refinement underlie the monocular bias of dLGN neurons. **a,** Scatter plot of mFD and AMPAR-based ODI (blue: contralateral dominated cells, red: ipsilaterally dominated cells; n = 91 cells). **b,** Probability of contralateral (blue) and ipsilateral (red) input dominance as a function of mFD, calculated from **a** by taking the mean eye preference in a range of +/− 0.2 mFD around each data point (n = 91 cells). **c,** Four example neighboring cell pairs with opposite eye preferences but similar mFD values. Traces from whole-cell voltage-clamp recordings at −70 mV (black) and +40 mV (gray) in response to double pulse light stimulation (durations for red and blue light indicated as bars over). **d,** Same as **a** but for NMDAR-based ODI (n = 91 cells). **e,** Binocularity index of AMPAR response is lower than the NMDAR response (n = 101 cells, Wilcoxon signed-rank test, W = 73, p < 0.001). **f,** AMPAR to NMDAR response ratio of monocular cells, and dominant and the non-dominant eye response of cells with a detectable AMPAR responses to both eye inputs (monocular: n = 48 cells, binocular: n = 53; monocular vs. DE: Mann-Whitney U test, U = 2565, p = 0.43; DE vs. NDE: Wilcoxon signed-rank test, W = 1377, p < 0.001). Image and morphology scale bar: 100 μm. Electrophysiology trace scale bars: 100 ms, 200 pA.

Importantly, NMDAR responses were on average less monocular than AMPAR responses (Fig. 7d,e). This was likely because the ratio of AMPAR to NMDAR responses were ~4x lower for the non-dominant eye compared to the dominant eye of the same cells, which were in turn indistinguishable from ratios of monocular cells (Fig. 7f). Furthermore, the AMPAR to NMDAR response ratios of both the dominant eye of binocular and monocular cells, were comparable to previously reported values in adult mice, whereas those of the non-dominate eye resembled ratios observed in juvenile animals^3,27^ This suggests that beyond synaptic input selection, synaptic refinement leads to further functional monocularity as not all of the synapses are functionally active (i.e. are “silent”, also see Fig. 2), or are functionally active but weaker. Thus, selection of specific inputs and their subsequent refinement are the primary mechanisms that lead to a strong functional monocular bias in dLGN neurons.

## Discussion

Our goal was to better understand eye-specific input selection in the mature mouse dLGN. We found that a small fraction of TC neurons shows binocular responses *in vivo*. In agreement with this, we showed *in vitro* that even though most dLGN neurons received detectable functional input from RGCs of both retinae, the majority of dLGN neurons were clearly dominated by input from one eye only. Eye-specific axo-dendritic apposition alone did not explain this prominent monocular bias. We rather find that functional input selection and refinement are the main factors leading to the general monocularity of dLGN neurons.

### Systematic assessment of mouse TC neuron binocularity *in vivo*

Recent anatomical and functional studies have challenged the long-standing dogma of strict eye-segregation in the adult mouse dLGN^2,9^. However, the degree and origin of binocularity remains debated, partially because of different recording techniques, different visual stimulus types, different analysis methods, and different TC neuron sampling^13^. We therefore evaluated the effect of these factors on *in vivo* TC neuron binocularity.

We found that a low fraction of TC neuron afferents (7%-21%, strongly dependent on the stringency of selection criteria) showed binocular responses with only minor and variable effects of stimulus type (grating or full-field luminance stimuli) and no effect of stimulus elevation. These numbers are still considerably lower than electrophysiological estimates but are in line with our previous observations^10^ and are similar to a recent report using a comparable recording approach in L1 of bV1^11^.

We observed small differences in TC neuron binocularity between cortical projection layers. Based on a previous study^31^, L1 recordings were expected to show a larger contribution of shell TC neurons in comparison to recordings from TC afferents in L4. This should have led to a lower fraction of ipsilaterally dominated and binocular axons, as our *in vitro* data clearly supports the hypothesis that the rodent dLGN shell is exclusively contralaterally dominated^22^. We indeed found a larger degree of TC neuron binocularity in L4 vs. L1. However, also TC neuron boutons in L1 clearly showed ipsilateral and binocular responses. It is possible, that L1-projecting TC neuron binocularity is partially driven by binocular input from the SC, which selectively innervates the dLGN shell^36^, but it remains to be further investigated to which degree the dLGN shell directly innervates L1 of bV1.

Given the non-linear nature of GECIs, our calcium imaging approach may underestimate small responses and, depending on distance to saturation regime, either emphasize or blunt strong responses^37^. We therefore cannot fully exclude that the far lower degree of binocularity in calcium imaging studies^10,11^ in comparison to electrophysiological reports^9,29^ is to some extent due to underestimation of weak, non-dominant eye responses. However, the relative similarity of our estimates across stimulus type, TC projection layer, and visuotopy suggests that technical differences between the recording modalities and analysis methods (e.g. source separation, smallest detectable response amplitude, threshold criteria used) play a larger role in the different estimates of binocularity than differences in stimulation and thalamic sampling. It is difficult to assess the exact nature of this discrepancy, but our *in vitro* mapping result of prominent functional dLGN neuron monocularity clearly shows that retinothalamic single cell convergence is unlikely to account for the large fraction of binocular responses observed in *in vivo* electrophysiological studies.

### Widespread binocular retinogeniculate convergence but limited functional binocularity

Previous work using retrograde mono-synaptic rabies tracing from the medio-dorsal tip of dLGN to the retinae has shown that roughly half (40%) of all dLGN neurons receive anatomical input from both eyes^2^. Our functional input mapping data is well in line with this finding. More than half of all dLGN neurons for which we mapped eye-specific retinogeniculate input strength with dual-channel CRACM received weak but clearly detectable input from the non-dominant eye (64%, based on NMDAR-mediated responses). In contrast to the aforementioned tracing study^2^ and previous *in vivo* data^9^ we also found a small number of neurons receiving exclusive input from the ipsilateral eye, corroborating our current and previous^10^ *in vivo* findings.

It is unlikely, that the limited degree of functional binocular retinogeniculate convergence that we observed *in vitro* is driving widespread binocular responses *in vivo* as observed in previous electrophysiological studies^9,29^. Non-dominant eye synaptic responses were far smaller (36-fold) than dominant eye responses and we were never able to elicit action potentials by non-dominant eye stimulation alone under resting conditions. Functional *in vitro* retinogeniculate binocularity was well comparable to functional *in vivo* binocularity within the sampled areas.

Our finding also fits well with recent electrophysiological *in vitro* and *in vivo* data on monocular retinogeniculate convergence^4,7^. Using a single-channel optogenetic mapping approach, a recent study^4^ found that the aggregate response to contralateral RGC axon photostimulation is dominated by only very few synapses. Our data extend this finding by showing that this mechanism of disproportionate synaptic strengthening at the retinogeniculate synapse is highly input-specific. Even with near-identical dendritic access to axons from both eyes, dLGN neurons select one input over the other in an almost all-or-none fashion.

### Morphological and functional mechanisms of eye-specificity

Our *in vitro* mapping data allowed us to assess the morphological origin of the prominent functional monocular bias of dLGN neurons. Throughout dLGN, we found that neurons were effectively functionally monocular and that binocular visual space was represented by a mixed population of largely monocular neurons. Importantly, neither eye-specific axonal segregation (1), nor dendritic orientation (2) alone were sufficient to explain this monocular bias:

1. Even when assuming the extreme case of completely non-overlapping, i.e. ‘binary’, eye-specific innervation of dLGN, simulated axo-dendritic overlap yielded a much higher estimate of retinogeniculate binocularity in comparison to our functional *in vitro* data, dismissing axonal segregation as the sole cause for functional monocularity. It should be noted that RGC inputs onto dLGN neurons are mostly located on proximal dendrites^5,8,38^. The majority (75%) of the radial mask weight used to calculate the fluorescence difference (rFD) lies under 70 μm from the soma. We therefore think that our estimates roughly match the effective reach of dLGN neuron dendrites with respect to retinogeniculate innervation. However, since the position of RGC inputs across the dendritic arbor as a function of distance from soma has not been quantified in detail, we were unable to integrate a more precise RGC synapse distribution into our calculations.
2. The orientation of cortical dendrites frequently has been shown to respect functional divisions resulting from segregated feedforward projection architecture. In V1 of cats and monkeys, dendritic fields of L4 stellate cells in the vicinity of eye-specific thalamocortical domains are asymmetric and favor arborizations in the ocular dominance column of their parent soma^39,40^. Similar biases in dendritic reach have been reported for other functionally defined compartments like cytochrome-C oxidase blobs in macaques^41^ (but see^42^) and in the barrel cortex of rats and mice^43^. Selective dendritic elongation and asymmetry could, in principle, increase or decrease sampling of one eye-specific RGC input over the other also in mouse dLGN. However, similar to cats^44^, primates^45^, and other reports in mice^30^, we found that dendritic arbors of dLGN neurons frequently crossed the diffuse border between the contra- and ipsilateral RGC axon projection zones. Furthermore, despite the frequent orthogonal orientation of dendritic elongation to the axon innervation gradient, this also did not lead to an increase of the average monocularity of dLGN neurons. Rather, even nearby cells with very similar access to RGC axon arbors from both eyes showed opposing eye preference.

### Input selection and synaptic refinement as mechanisms for functional dLGN monocularity

We functionally identified inputs via eye-specific stimulation, which allowed us to assess a range of parameters of retinogeniculate transmission with input-specificity. We found that even though most dLGN neurons tested received input from both eyes, the fine-scale synaptic properties clearly favored monocularity. Not only was the synaptic response of the non-dominant eye far weaker in total for both AMPAR- and NMDAR-mediated responses, but we also found that the non-dominant eye showed a significantly lower AMPAR / NMDAR ratio – to the degree of full AMPAR-silencing in some non-dominant eye connections. The non-dominant eye therefore is hardly able to contribute to dLGN neuron binocularity, because of both absolute synaptic strength and partial or complete synaptic silencing. We estimate that of ~20,000 dLGN neurons in total^46^, only ~200 cells would receive roughly equal input from both eyes. It remains to be tested if these few cells fulfil a specific role in binocular vision.

The lower AMPAR / NMDAR ratio of the non-dominant eye in the adult mouse dLGN is reminiscent of the situation at pre-eye-opening retinogeniculate synapses. It has been shown frequently that the contribution of AMPAR over NMDAR ratio increases prominently during development^3,27^, alongside the strengthening of a small number of dominating inputs^3,4^. Fully or partially silent non-dominant eye synapses therefore bear signs of a perinatal retinogeniculate developmental process. The suppression of non-dominant synapses and the residual binocularity picked up by rabies tracing could be interpreted as the result of a protracted process of functional eye-specific synaptic refinement as has previously been suggested for the functional silencing of convergent monocular RGC input^4^. To disentangle the dynamics of input-specific developmental changes, further research would be needed to understand at which point in development non-dominant eye inputs are functionally silenced and if functional refinement may progress even further beyond the ages tested in this study. Whether or not also monocular retinogeniculate convergence of different RGC types^4,7^ is limited by silencing of non-dominant inputs remains to be established as well. The dual-channel approach we present here together with genetic access to different RGC types^47^ should now make this question tractable.

Functionally, the retinogeniculate synapses containing a larger fraction of NMDARs could serve as a silent pool of synapses that may become rapidly recruited through activity-dependent insertion of AMPARs. Whether or not residual functional binocularity plays a role in experience-dependent plasticity by un-silencing of non-dominant eye input could be the subject of future studies. However, without major changes in absolute synaptic strength we consider it unlikely that un-silencing alone would be able to explain the shifts in eye-specific responsiveness of LGN cells we^10^ and others^29^ have observed after both adult and juvenile ocular dominance plasticity.

In summary, we conclude that fine-scale input selection and synaptic refinement, partially via selective glutamate receptor allocation, is the dominant factor in determining limited functional binocular retinogeniculate convergence onto single dLGN neurons. Even though mouse dLGN clearly shows a large degree of eye-specific structural convergence, we find that mature mouse dLGN is effectively functionally monocular. The apparent mismatch between previous structural and functional data on dLGN binocularity can therefore be explained by our data. Whether or not non-dominant inputs can be rendered functional by experience-dependent plasticity^48^ or if they are further curbed in a process of protracted adult refinement will be interesting questions for future research.

## Material and Methods

### Animals

All experimental procedures were carried out in compliance with institutional guidelines of the Max Planck Society and the local government (Regierung von Oberbayern). Female wild type C57bl/6 mice as well as male and female Scnn1a-Tg3-Cre mice were used. Mice were housed under a 12 hour light-dark cycle with food and water available ad libitum.

Craniotomy, virus injections and head plate implantation were performed at P45-55 on Scnn1a-Tg3-Cre mice. Animals were usually group housed. After cranial window and head plate implantation animals were singly housed. All intravitreal eye injections were performed on P30-P48 old female C57bl/6 mice. *In vitro* brain slice experiments were performed at P67-P108.

### Virus design and production for *in vivo* experiments

Two recombinant adeno-associated viruses (AAVs) encoding Flpo and jGCaMP7b were created to control transfection sparseness in the Cre-expressing Scnn1a-Tg3-Cre mouse line. The pAAV1-EF1α-F-Flex-jGCaMP7b-WPRE viral construct was generated by subcloning the jGCaMP7b coding sequence from pGP-AAV-Syn-jGCaMP7b-WPRE (Addgene plasmid number #104493) into pAAV-EF1α-F-Flex-Kir2.1-T2A-tdTomato-WPRE (Addgene plasmid number 60661), replacing Kir2.1-T2A-tdTomato using HindIII for the donor and BsrGI for the target vector, which were blunted and restricted using BamHI. Correct insertion was verified by sequencing and control restrictions using HpaI/BamHI and HpaI/EcoRV. The pAAV2/1-CAG-Flex-Flpo-WPRE viral construct was generated by subcloning the Flpo coding sequence from pAAV-hSyn-Flpo-WPRE (Addgene plasmid number 60663) into pAAV-CAG-Flex-ArchT-tdTomato-WPRE (Addgene plasmid number 20305), replacing ArchT-tdTomato using EcoRI-HF for the donor and KpnI-HF and XboI for the target vector, which were blunted. Correct insertion was verified by control restrictions NheI/HindII and NheI/NotI. Both vectors were packaged into AAVs by the vector core of the University of Pennsylvania Gene Therapy Program (pAAV1-EF1α-F-Flex-jGCaMP7b-WPRE: 1.38 x 10^13^ genome copies/ml, AAV2/1-CAG-Flex-Flpo-WPRE: 1.244 x 10^13^ genome copies/ml).

### Solutions

Cortex buffer for *in vivo* surgeries contained (in mM): 125 NaCl, 5 KCl, 10 glucose, 10 HEPES, 2 CaCl2 and 2 MgSO_4_. The buffer was sterilized and maintained at pH 7.4.

The cutting solution for *in vitro* experiments contained (in mM): 85 NaCl, 75 sucrose, 2.5 KCl, 24 glucose, 1.25 NaH_2_PO_4_, 4 MgCl_2_, 0.5 CaCl_2_ and 24 NaHCO_3_, 310-325 mOsm, bubbled with 95% (vol/vol) O_2_, 5% (vol/vol) CO_2_. Artificial cerebrospinal fluid (ACSF) contained (in mM): 127 NaCl, 2.5 KCl, 26 NaHCO_3_, 2 CaCl_2_, 2 MgCl_2_, 1.25 NaH_2_PO_4_ and 10 glucose, 305-315 mOsm, bubbled with 95% (vol/vol) O_2_, 5% (vol/vol) CO_2_. Cesium-based internal solution contained (in mM): 122 CsMeSO_4_, 4 MgCl_2_, 10 HEPES, 4 Na-ATP, 0.4 Na-GTP, 3 Na-L-ascorbate, 10 Na-phosphocreatine, 0.2 EGTA, 5 QX-314, and 0.03 Alexa-594, pH 7.25, 295-300 mOsm. K-gluconate-based intracellular recording solution contained (in mM): 126 K-gluconate, 4 KCl, 10 HEPES, 4 Mg-ATP, 0.3 Na-GTP, 10 Na-phosphocreatine, 0.3-0.5% (wt/vol) Neurobiotin tracer and 0.03 mM Alexa-594, pH 7.25, 295-300 mOsm.

For brain slice clearing, the blocking buffer contained: 10% Normal Goat Serum, 2% Triton X-100, 0.2% Sodium Azide in PBS. The antibody buffer contained: 1% Normal Goat Serum, 0.2% Triton X100, 0.2% Sodium Azide in PBS. The washing buffer contained: 3% NaCl, 0.2% Triton X-100 in PBS. The permeabilization buffer contained: 2% Triton X-100 in PBS.

### Thalamic virus injection and chronic-window preparation

To reduce the cortical density of TC afferents (Fig. 1b) and thereby minimize potential contamination of bouton calcium signals by neighboring neurites in densely thalamorecipient L4 of bV1^49^ a method for sparse double-conditional transduction was devised in a mouse line with thalamic specificity^50^. Scnn1a-Tg3-Cre mice were anesthetized via an intraperitoneal injection of a mixture of fentanyl (0.05 mg/kg), medetomidine (0.5 mg/kg) and midazolam (5 mg/kg). Analgesia was supplied both locally via topical application of 10% lidocaine and globally via subcutaneous injection of carprofen (5 mg/kg before surgery, on the first and second day of post-surgical recovery). Mice were placed on a heated blanket (37°C; Homeothermic blanket with rectal probe) to ensure thermal homeostasis during the procedure. Eyes were protected from dehydration by applying eye cream to both eyes. Wound sepsis was prevented by application of an iodine solution on top of the skin before the first incision. Mice were fixed into a motorized stereotaxic (StereoDrive Motorized Stereotaxic. Neurostar, Tuebingen, Germany) and a circular craniotomy (4mm diameter) was drilled) on top of primary visual cortex of the right hemisphere. Dehydration of the brain was prevented by periodic application of cortex buffer on top of the brain. Sparse dLGN transduction was achieved by a targeted stereotactic injection (2.06 mm anterior, 2.05 mm lateral relative to bregma) through a glass syringe (NanoFil needles, beveled, gauge 36. NanoFil 10 μl syringe. World Precision Instruments). 50-70 nl of a mixture of AAV2/1-CAG-Flex-Flpo-WPRE (titer: 2.488 x 1010 genome copies/ml) and AAV1-EF1α-F-Flex-jGCaMP7b-WPRE (titer: 1.104 x 1010 genome copies/ml) were injected at a depth of 2.80-2.85 mm and a rate of 0.5 nl/s using an automated microinjector (Neurostar). The craniotomy was sealed by a glass cover slip (4mm diameter) and super glue. A custom machined head-bar was attached to the skull using a mixture of dental cement and pigment.

After surgery, anesthesia was counteracted by a subcutaneous injection of a mixture of Naloxone (1.2 mg/kg), Flumazenil (0.5 mg/kg) and Atipamezol (2.5 mg/kg). Expression of GECI at the level of bV1 could be observed 4-5 weeks after surgery.

### Intravitreal eye injections

Intravitreal eye injections were performed on mice that were intraperitoneally anesthetized with a mixture of fentanyl (0.05mg/kg), midazolam (5mg/kg) and medetomidine (0.5mg/kg). 5 mg/kg carprofen was administered as an analgesic on the day of surgery and on the following two days. Before starting the surgery, all surgical instruments were heat-sterilized and washed with ethanol. 5 μl Hamilton syringes (Model 75 RN SYR, Hamilton) tipped with a 32G blunt needle (Small hub RN needle, Hamilton) were rinsed several times with distilled H_2_O, then ethanol and again with distilled H_2_O. Chronos-(AAV2/2.Syn-Chronos.EGFP) and ChrimsonR-expressing AAVs (AAV2/2.Syn-ChrimsonR.tdT) were loaded into separate Hamilton syringes, and any air was expelled by repeated ejection and loading of the virus. The animal was then fixated in a stereotax and the eye that was not injected first was kept damp with eye drops (Oculotec, Novartis). The target eye was held in place using forceps by the connective tissue at the back of the eye and a small hole was punctured just behind the corneo-scleral junction using a 0.4 mm sharp syringe tip. A virus loaded Hamilton syringe, controlled using a micromanipulator (M3301R, WPI), was then inserted into this opening at an oblique angle to avoid damaging the lens and any vitreous fluid that extruded from the eye was removed using a cellulose sponge. 1-2 μl of virus was injected into the eye and the syringe left in place for 4 min to allow the virus to disperse. After removing the Hamilton syringe the eye was covered with eye cream and the procedure was repeated with the other eye using the second virus. Viruses expressed for five to nine weeks. After sacrificing for *in vitro* experiments, animals were periodically checked for lens damage (cataracts), of which none were found using this protocol.

### *In vivo* intrinsic optical signal imaging

Reflected-light imaging of intrinsic optical signals as a readout of ocular dominance and retinotopy was performed as described previously^10,51^ under anesthesia (Initial anesthesia: one intraperitoneal injection of fentanyl (0.035 mg/kg), midazolam (3.5 mg/kg) and medetomidine (0.35 mg/kg); anesthesia during imaging was maintained by hourly injection of a lower dose (25% of induction level) of the anesthetic mixture). In brief, the optical axis of the camera was aligned to be orthogonal to the cranial window in each mouse. The surface of the brain was first side-illuminated with light from a 530 nm LED to visualize the blood vessel pattern and subsequently from two sides with a 735 nm LED for intrinsic signal imaging. Signals were band-pass filtered at 700/40 nm and collected with a CCD camera (12-bit, 260 x 348 pixels, 40 Hz for localization of primary visual cortex using drifting square wave gratings; 12-bit, 512 x 512 pixels, 15 Hz for mapping primary visual cortex retinotopy using drifting reversing checkerboard bars) through a 4x air objective (NA 0.28). The imaging plane was set to be 400-450 μm below the pial surface. Acquisition and analysis software were custom written in MATLAB.

### *In vivo* two-photon imaging

*In vivo* two-photon Ca^2+^ imaging was performed as described earlier^10,52^. Briefly, jGCaMP7b was excited by a 80 MHz pulsed femtosecond Ti:sapphire laser (MaiTai eHP, Spectra-Physics) tuned to 940 nm. Fluorescence signals were detected by a 16x NA 0.8 objective using GaAsP photomultiplier tubes (short-pass filtered at 720 nm and band-pass filtered at 525/50 nm). Fields-of-views of 64×50 μm were acquired at a resolution of 512×512 pixels at 30.4 Hz by bidirectional scanning using an 8KHz resonant scanner and a Pockels cell for beam turnaround blanking. For volumetric multiplane imaging the objective was rapidly moved in the z axis by a high-load piezoelectric z scanner, yielding in a pseudo-simultaneous acquisition of 4 subsequent inclined planes either 15 or 10 μm apart at an effective frame rate of 7.6 Hz. For imaging in layer 1, volumes were taken between 10 and 100 μm and in layer 4 between 300 and 450 μm below the pial surface. The average laser power under the objective was kept at ≤ 50 mW during imaging. A commercial rotating body system (Bergamo II, Thorlabs) was used to align the optical axis orthogonally to the imaging window. The data was acquired using ScanImage 4.2^53^ and custom-written hardware drivers.

*In vivo* two-photon imaging experiments were performed under light anesthesia and controlled thermal homeostasis (mice placed on a heated blanket). Initial anesthesia was achieved by one intraperitoneal injection of fentanyl (0.035 mg/kg), midazolam (3.5 mg/kg) and medetomidine (0.35 mg/kg), while anesthesia during imaging was maintained by hourly injection of a lower dose (25% of induction level) of the anesthetic mixture.

### Visual stimulation

Visual stimulation was performed as described previously^10,52^. Briefly, visual stimuli were generated using Matlab and the Psychophysics Toolbox^54,55^. Visual stimuli were presented on a gamma-corrected liquid crystal display centered in the visual field of each mouse.

Ocular dominance was measured by either drifting grating or full-field luminance stimulation. Drifting square wave gratings (12 directions, spatial frequency: 0.04 cycles/degree; temporal frequency: 3 cycles/s) were presented in pseudorandom order to the right eye, the left eye, and both eyes using motorized eye shutters (6 trials per condition). Stimuli were presented in the binocular visual field of the mouse (−15° to 35° elevation, −25° to 25° azimuth relative to midline) for 5 seconds on a gray background, followed by 6 seconds of full-field gray at 50% contrast. Full-field luminance stimuli were adapted from^7^ and consisted of a three second long step-wise full-contrast change followed by two sinusoidal light intensity modulations with 8 seconds of increasing frequency and 8 seconds of increasing contrast on a gray background with an inter-trial interval durations of 2 seconds at uniform gray. The initial 9 seconds of full-contrast change are denoted as ‘flash’ and the complete luminance stimulus as ‘full-chirp’. Luminance stimuli were again presented in pseudorandom order to the right eye, the left eye and both eyes (10 trials per condition). To control for potential light-leakage we also presented luminance stimuli when both eyes were covered by the motorized eye shutters (Supplementary Fig. 1).

For intrinsic optical signal imaging of binocular primary visual cortex, two different stimuli were used. First, drifting square wave gratings (spatial frequency: 0.04 cycles/degree; temporal frequency: 2 cycles/s) were presented as two patches (20×40 visual degrees) randomly at two adjacent positions (interface centered in azimuth). Stimuli were presented for 7s (eight directions, changing every 0.6s) to both contralateral and ipsilateral eye using motorized eye shutters (7 trials per condition). Inter-trial intervals consisted of 8s of a full-field gray screen^10,52^. Second, a drifting bar consisting of a reversing checkerboard pattern (special frequency: 0.04 cycles/degree; temporal frequency: 2 Hz) was periodically swept in both cardinal axes over a gray background (speed: 20 degrees/s, 10 trials per stimulus condition per eye) to map the retinotopy of primary visual cortex as described previously^56^.

### Acute brain slice preparation

Acute brain slices were obtained on the day of *in vitro* experiments, as previously described^57^. In brief, mice were deeply anesthetized with Isoflurane in a sealed container (>100 mg/kg) and rapidly decapitated. Coronal sections of dLGN (320 μm) were cut in ice cold carbogenated cutting solution using a vibratome (VT1200S, Leica). Slices were incubated in cutting solution in a submerged chamber at 34°C for at least 45 min and then transferred to ACSF in a light-shielded submerged chamber at room temperature (21°C) until used for recordings. The expression pattern of Chronos-EGFP and ChrimsonR-tdTomato within the dLGN was screened using fluorescence detection googles (Miners lamp, BLS Biological Laboratory Equipment) with different excitation light and filters (green and red) during the slice preparation. Only slices with visibly sufficient transduction of Chronos-EGFP and ChrimsonR-tdTomato were considered for experiments. Brain slices were used for 6-12 hours.

### *In vitro* electrophysiological recordings

Brain slices were mounted on poly-D-lysine coated coverslips (10 mm) and then transferred to the slice perfusion chamber and visualized with an upright microscope using infrared differential interference contrast optics (iDIC). Whole-cell voltage- and current-clamp recordings of thalamic neurons were performed at room temperature in ACSF with borosilicate glass patch pipettes (resistance of 4-5 MΩ) filled with Cs- and K-gluconate-based internal solution, respectively. Series resistance was usually below 30 MΩ. Data were acquired with Multiclamp 700 B amplifiers (Axon instruments). Voltage clamp recordings were filtered at 4-8 kHz and digitized at 10-20 kHz. Experiments were performed in the presence of the GABAA receptor antagonist bicuculline (20 μM) to block inhibitory di-synaptic connections.

### Dual-color sequential photostimulation

Dual-color photostimulation experiments were performed at two identical setups. At the beginning of the experiment, an overview two-photon image stack of the dLGN was obtained using λ = 940 nm to simultaneously excite EGFP (emission filter: 525/50-25, Semrock) and tdTomato (emission filter: 607/70-25, Semrock) using a low magnification objective (i.e. 16x, 20x for the two setups). This overview was used as an orientation tool in between electrophysiological recordings, when selecting a new target cell within a region of interest.

For dual-color photostimulation, a single full-field laser pulse was used in order to map the net synaptic input to the recorded cell. 475 and 673 nm illumination was provided by a blue and red laser source (473 and 637 nm, S3FC473, S4FC637, Thorlabs). The laser light from the two laser sources were combined using a 2-color combining fiber (RB42F1, Thorlabs) before entering the microscope head. The combined fiber was then plugged into the microscope head using a fiber-collimator (F230FC-A, PAF-X-15-A, Thorlabs). Galvanic scan-mirrors controlled the positioning of the laser beam and the scan and tube lens were arranged to under-fill the back aperture of the microscope objective (40x LUMPlan, 0.8 NA Olympus or 4x Plan N 0.1 NA Olympus), resulting in a pencil-shaped beam with a diameter of 2.35-2.6 mm.

To adequately measure a cell’s maximal input evoked by blue and red light stimulation we used two step protocols, jointly called the maximal current response protocol: First, the blue laser intensity was increased and the maximal evoked current was obtained for each dLGN neuron (11 steps, irradiance range: 0-3 mW/mm^2^, Fig. 2c, top). Each blue pulse stimulus was separated by 10 s. Subsequently, we used sequential photostimulation (a red 250 ms long stimulus immediately followed by a 50 ms long blue light stimulus^33^) with increasing red laser intensity to obtain the maximal current response evoked by the other eye in the same dLGN neuron. The blue laser intensity that evoked response saturation in the first step protocol was used as a constant paired stimulus for the second step protocol (11 steps, irradiance range: 0-5.1 mW/iW, Fig. 1F, bottom). Only, the second, cross-talk-eliminating stimulation protocol was used to obtain relative eye-specific input strength. Both protocols were applied at −70 and +40 mV membrane holding potential to capture evoked AMPAR- and NMDAR-mediated responses in the same cell.

In a subset of cells, current-clamp experiments were performed to probe under which conditions dominant and non-dominant eye inputs could evoke action potentials. For this, three steps were taken: First, ocular input type and dominance was established by means of the above outlined maximal current response protocol in voltage-clamp. Second, intrinsic properties (e.g. spiking threshold) were determined by applying hyper- and depolarizing current pulses after switching to whole-cell current-clamp. Lastly, the established power levels of the first step, eliciting maximal responses were taken and five iterations of blue- and red light sequential photostimulation were performed. This was first done at resting potential, and subsequently current was injected, depolarizing the cell and step-wise (10 mV steps) bringing it closer to the spiking threshold, established in the second step, until either the spiking threshold was reached, or action potentials were elicited by both stimulation colors (Fig. 2h, 2i).

In a subset of cells, intravitreal injections were only performed monocularly via ChrimsonR-expressing AAVs (AAV2/2.Syn-ChrimsonR.tdT) to explore potential crosstalk effects of blue light stimulation (Fig. 2b). To determine the intensity ratio between red and blue light for successful crosstalk prevention, the maximal current response protocol was used (see above, Supplementary Fig. 2a). To determine the duration of the red pre-stimulation which successfully prevents the consequent blue light stimulation from eliciting a cross-talk-driven response, maximal response intensities (as determined during the maximal current response protocol) were taken with a variable red light stimulation window (1 ms, 10 ms, 250 ms.) and a fixed blue light stimulation window (50 ms, Supplementary Fig. 2b, c).

To quantify the effect of red light-induced ChrimsonR desensitization, we used maximal response intensities (as determined during the maximal current response protocol) and first used only red light stimulation three times with a 5 s ISI, then performed sequential photostimulation (red and blue) also three times with the same ISI (Supplementary Fig. 2d, e). Repetitive light stimulation for 250 ms with single red pulses of 637 nm led to a pronounced rundown of peak postsynaptic current (PSC) amplitude in our preparation (Supplementary Fig. 2e). This rundown was prevented by pairing a 250 ms long red pulse with blue light stimulation at 473 nm (50 ms) as used for sequential photostimulation in our dataset (Supplementary Fig. 2e).

### Immunohistochemistry and brain slice clearing

After *in vitro* electrophysiological recordings, acute slices were stored in 4% PFA at 4°C until immunohistochemistry and brain clearing was performed: In short, the brain slices were washed (3×10 min in PBS) and then incubated in permeabilization buffer overnight at 4°C. The brain slices were incubated in blocking buffer for 8 hours at RT, then incubated in the primary antibody Rabbit Anti-Calbindin D28k (1:2,000 in Antibody buffer, Swant CB-38a) overnight at RT and subsequently at 4°C on a shaker for 3 days. After washing the brain slices overnight, the slices were incubated in the secondary antibody Anti-Rabbit Alexa-647 (1:200 in Antibody buffer, Thermo Fisher A21244) at 4°C on a shaker for 2 days. Slices were subsequently washed in washing buffer and then in PBS.

For brain slice clearing, slices were incubated in RapiClear 1.47 *(SunJin Lab Co.) and cleared for approximately 3 hours. Slices were then embedded in RapiClear solution and covered with a coverslip on a 300 μm spacer.

### Confocal imaging of fixed brain slices

Cleared slices were imaged using a confocal microscope (Sp8, Leica) at a voxel size of 1.614 x 1.614 x 4.0 μm. Equipped with an argon-ion laser (used at 488 nm), as well as a diode pumped solid-state (DPPS) laser (561 nm) and a Helium-Neon laser (633 nm), we acquired three channels sequentially through a 20x HCX APO L 20x/1.00 WATER objective (Leica): Spectral detection windows were set to capture EGFP (493-555 nm) and Alexa-647 (staining against calbindin, 638-750 nm) – imaged simultaneously -, and tdTomato (565-628 nm). Scan speed was set to 600 Hz. Online averaging of three line scans resulted in the individual images, which were mosaic merged (10% overlap, LAS X, Leica) to create the final, tiled image of the entire part of the LGN that the slice contained.

### *In vivo* calcium imaging analysis

Data analysis was performed in Python (v3.7, distribution by Anaconda Inc., Austin, TX) using custom routines and published packages.

### Pre-processing

Regions of interest (ROI) to extract calcium signals of putative TC neuron boutons were automatically generated using suite2p (v0.71)^58^. The pipeline included non-rigid image registration followed by activity-based segmentation of regions of interest (ROIs). The segmentations were curated semi-automatically using anatomical criteria to, e.g., exclude ROIs that corresponded to larger patches of low contrast signal. Automatic curations were inspected and corrected manually using the suite2p GUI. All analysis was performed on the neuropil-corrected, detrended, and deconvolved raw fluorescence data^58^. Neuropil correction was performed using a circular annulus around each detected putative bouton (at least three-fold radius of the detected bouton radius, spaced 0.2 μm away from the ROI circumference), excluding overlap with other selected cells and neuropil bands. The neuropil-corrected average fluorescence per ROI was calculated as Fbouton_corrected = Fbouton_raw – r × Fneuropil with a correction factor r fixed to 0.7^52,59^. The neuropil-corrected traces were baseline-subtracted (rolling maximum of rolling minimum) and then deconvolved using the OASIS algorithm (decay time constant *τ* = 1 s)^58,60^. Deconvolved data was used to allow for better resolution of high-frequency-modulated changes during chirped full-field-luminance stimuli (Figure 1c).

### Data analysis

Responses to drifting grating stimuli were extracted from either trial-averaged or single-trial data as mean absolute change in the deconvolved fluorescence (R) over the full stimulus interval for each condition (ΔR = R – Rbase), with Rbase representing the mean deconvolved fluorescence response during 1.5 s of full-field gray (i.e. ‘blank’) stimulation preceding stimulus onset. To determine visual responsiveness to contra- and ipsilateral eye stimulation, a one-way ANOVA was performed over all single-trial R and Rbase ‘blank’ values per stimulus direction. Three different threshold criteria were used to assess responsiveness (p < 0.05, p < 0.01, and p < 0.001, Supplementary Fig. 1c). To assess monocular and binocular categorical responsiveness no positivity constraint was included so that both increases and decreases in activity were scored as significantly responsive (Fig. 1 d,e).

The ocular dominance index (ODI, Fig. 1f) was calculated only for positive responses of significantly responsive boutons and extracted from the trial-averaged preferred direction responses to monocular stimulation as

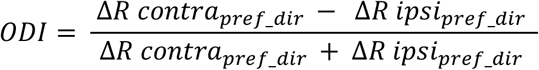

Non-significant responses were set to zero.

Modulation of activity during binocular stimulation compared to monocular stimulation was assessed for significant positive responses as the ratio of the preferred direction response to both-eye stimulation over the preferred direction response of the dominant eye during monocular stimulation:

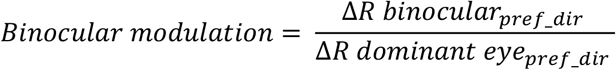

Responses to full-field luminance stimuli were analyzed separately for the initial ‘flash’ part and the complete luminance stimulus (‘full-chirp’, Fig. 1c). On- and Off-responses to the ‘flash’ part were extracted from either trial-averaged or single-trial data as mean change in the deconvolved fluorescence over the full stimulus interval both for the bright as for the dark step (ΔRbright = R – Rblack and ΔRdark = R – Rblack), with Rblack representing the mean pre-stimulus deconvolved fluorescence during the three seconds of full-field black stimulation preceding stimulus onset. To determine visual responsiveness to contralateral and ipsilateral eye stimulation, a one-way ANOVA was performed over the Rbright, Rdark, and Rblack values. Three different threshold criteria (denoted as ‘p-threshold’) were used to assess responsiveness (p < 0.05, p < 0.01, and p < 0.001, Supplementary Fig. 1c). As previously, no positivity constraint was included to assess categorical monocular responsiveness (Fig. 1c,e).

Ocular dominance in response to ‘flash’ stimulation (Fig. 1f) was calculated similar as for gratings but in this case using the mean response of the preferred stimulus polarity (i.e. either On- or Off-response). Non-significant responses were set to zero.

Responsiveness of individual boutons to the ‘full-chirp’ stimulus was assessed by separating the eye-specific stimulus responses into two segments of equal length and computing the average between-trial correlations within-segment and across segments^7^. Responsiveness was scored using a Wilcoxon rank-sum test of within-segment correlations vs. across-segment correlations (three different threshold criteria p < 0.05, p < 0.01, and p < 0.001, no positivity constraint, Fig. 1c,e, Supplementary Fig. 1).

To concisely display the full variety of stimulus-evoked activity of our dataset in a meaningful way, the stimulus-aligned data (concatenated over monocular and binocular stimulation) was z-scored and then sorted separately for significantly contralateral, ipsilateral, and binocular (contralateral and ipsilateral) boutons. For this we used a manifold embedding algorithm (‘Rastermap’, www.github.com/MouseLand/RasterMap)^61^ so that boutons with correlated activity lie next to each other (Fig. 1c,d, Supplementary Fig. 1a,b).

For comparison of eye-specific partitioning across experimental categories and statistical thresholds, multivariate loglinear models were used^62^. Briefly, we generated a nested contingency table with the response categories ‘stimulus (three level), ‘elevation’ (two level), ‘layer’ (two level), ‘threshold-p’ (three level), and ‘eye-specific response fraction’ (three level). We were interested in whether a loglinear model including select pairwise interactions (e.g. for eye-specific response fraction X Stimulus: log(frequency) = log(eye-specific response fraction) + (log(eye-specific response fraction) x log(stimulus)) + log(elevation) + log(layer) + log(threshold-p)) performed better than the noninteraction main model (e.g. log(frequency) = log(eye-specific response fraction) + log(stimulus) + log(elevation) + log(layer) + log(threshold-p)). For model comparisons the log likelihood-ratios were compared using chi-squared statistics. Statistical analysis was performed in R.

To assess the dispersion of binocularity estimates across conditions the mean absolute deviation (MAD) around the mean binocularity estimate for a single responsiveness threshold p-value is reported.

### Dual-color sequential photostimulation analysis

For quantification of light-evoked PSC responses, electrophysiological traces were first baseline adjusted. This was performed by measuring the average current amplitude within a temporal window of 100 ms prior to red stimulation and subtracting this from all traces.

To determine light-evoked responsiveness, a nonparametric signed-rank test was performed over the average current during the response and baseline windows for the respective laser stimulation for the last 6 steps of the 11-step stimulation protocol. Current deflections with p-values < 0.05 were identified as significant light-evoked PSCs.

The response window was 27 ms for AMPAR and 104 ms for NMDAR responses and was measured 3 ms after the start of the respective laser stimulation. The baseline window was of similar length compared to the response window and was measured prior to the start of the respective laser stimulation. Since the NMDAR response evoked by the second laser stimulation (blue) was overriding on the decay phase of the response to the first stimulation (red), an exponential fit to the decay phase of the first response (single or double exponential, whichever had a higher df adjusted R^2^) was subtracted from the trace before quantifying the baseline and response periods of the second response.

Ocular dominance *in vitro* was determined by the ocular dominance index (ODI) for each dLGN neuron for AMPAR and NMDAR responses separately:

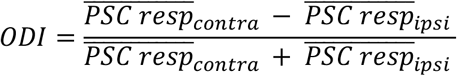

where PSC_resp_ was determined in the following way: 1) if the cell received significant input evoked by red and blue laser stimulation, the maximum among all baseline subtracted peak responses of the 11 steps was used. 2) if the cell received input evoked by blue laser stimulation only, the average of the peak responses for the last 6 steps of the 11-step stimulation protocol was used. PSC_resp_ was set to zero for unresponsive cells.

Similarly, the crosstalk suppression index (Supplementary Fig. 2c) was calculated as:

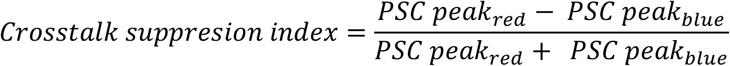

where PSC peaks were determined as responses with a deflection higher than 3 S.D. over the baseline (baseline for red stimulation: 100 ms prior to stimulation; for blue stimulation: 100, 2, 1 ms for red stimulation durations of 250, 10 and 1 ms).

### Morphology reconstruction and analysis

After successful recording and filling with Alexa-594 a detailed structural two-photon image stack of the dendritic morphology of the entire cell was acquired with excitation light of λ = 810 nm (emission filter: 607/70-25, Semrock, Rochester, USA) using ScanImage 4.2^53^. The structural image stacks typically consisted of ~100 sections (1024 x 1024 pixels; 0.3-0.8 μm per pixel) collected in z-steps of 1.2 μm. Subsequently, a second identical image stack was acquired at λ = 940 nm to locate the cell with respect to the RGC axonal fibers expressing tdTomato and EGFP.

Reconstruction of dendritic cell morphologies was performed manually using the Simple Neurite Tracer of ImageJ^63^. Reconstructions were quantitatively analyzed with custom-written MATLAB code and the open-source TREES toolbox^64^. The soma position was defined as the mean position of all soma nodes (indicated points along a dendrite, axon or soma). Dendritic reach was defined as the maximum Euclidean distance of all dendritic nodes from the soma and total dendritic length was measured as the sum of all dendritic internode sections.

For Sholl analysis, the number of intersections between dendrites and concentric spheres centered on the soma was determined at increasing distances from the soma (20 μm increments) and maximum Sholl crossing was then defined as the sphere distance with the most dendrite crossings. Dendritic orientation index (DOi) was calculated as defined in^30^. In short, axial planes were defined by rotating (in the coronal plane) 4 equal quadrants around the cell until the ratio of dendritic crossings, as defined in Sholl analysis using 20 μm spheres, in one quadrant compared to its opposing quadrant was maximized. These two quadrants were defined as a1 and a2 (combined defined as axial plane a) while the other two opposing quadrants were defined as b1 and b2 (axial plane b). DOi was then calculated by dividing the number of crossings in the axial plane with most crossings (either a or b) by the number of crossings in the orthogonal axial plane.

For all further analysis the dendritic morphology traces were interpolated in 3D (to produce interpolated nodes of equal spacing instead of the manually annotated nodes) and oriented to a common coordinate system. To align morphologies to the confocal image stacks, the overview two-photon image stack of dLGN was aligned, via manual rotation (in 10° increments) and reflection along the x or y axis, to best match the orientation of the dLGN confocal image stack of each brain slice. This transformation was then simply applied to the interpolated morphologies. These aligned morphologies were used to calculate the morphology-based FD (mFD; see below). To compare morphological orientation across brain slices, the confocal image stacks of all dLGNs were rotated manually, using custom written MATLAB code, to match the right dLGN of the Allen common coordinate framework (ACCF)^65^. These transformations were subsequently applied to the dendritic morphologies. Both left and right dLGNs were aligned to the right hemisphere of the ACCF. The left to right axis was therefore referred to as the medial to lateral axis. The direction of an angle was defined to be within the range of −180° to 180 and the orientation of an angle as within the range of – 90° and 90°. A vector of magnitude = 1 and direction = −90° in this coordinate system corresponds to 1μm displacement from ventral to dorsal, while a vector with magnitude 1 and direction = −180° corresponds to 1μm displacement from lateral to medial.

Dendritic asymmetry direction (*Asym_dir_*) and orientation (*Asym_ori_*) were defined as:

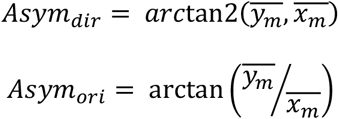

Where 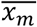 is the mean of all interpolated dendritic nodes in the medio-lateral axis and, 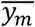 is the mean of all interpolated dendritic nodes in the ventro-dorsal axis. The absolute normalized difference between the number of nodes on either side of an orthogonal line to the asymmetry orientation and intersecting the soma, was then defined as the asymmetry magnitude (*Asym_mag_*; ranging from 0 to1):

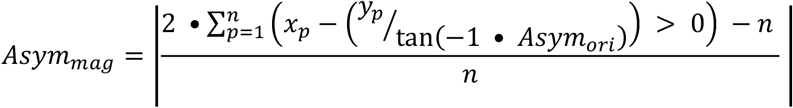

Where *x_p_* and *y_p_* are the coordinates of each node, and n the number of nodes.

Dendritic elongation was calculated by first performing principal component analysis (PCA) on the x and y components of the interpolated dendritic morphology of a cell. Elongation orientation (*Elong_ori_*) was then defined as:

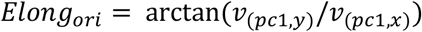

Where *v*_(*pc1, x*)_ and *v*_(*pc1, y*)_ are the x and y coefficients of the eigenvector of the first principle component (PC). Elongation magnitude (*Elong_mag_* ranging from 0 to 1) was defined as:

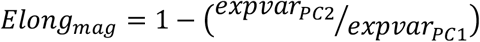

Where *expvar_PC1_* and *expvar_PC2_* are the explained variances of PC1 and PC2. This measure was partially derived from Morgan et al., 2017.

To determine if the dLGN morphologies can be clustered into discrete groups, PCA was performed on all six quantitative morphology measures: dendritic reach, total dendritic length, maximum Sholl crossing, DOi, asymmetry magnitude and elongation magnitude. All morphologies from cells in the dLGN, which did not have a distinct inhibitory type morphology and passed the quality control criteria, were included in this analysis even if no functional data was available. Hartigan’s dip test^66^ (http://www.nicprice.net/diptest/) for multimodality was then performed on the distribution of coefficients of the first two principle components.

### Re-identification of dLGN cells and fluorescence difference stack generation

After acquisition of the confocal image stack of each cleared brain slice ImageJ was used to first manually identify the position of each cell, either by re-identifying the cell itself (if the cell body remained adequately filled with Alexa-594 after staining and clearing) or by the structure of local RGC axons. The dLGN was also manually traced at multiple z positions to produce a convex hull of dLGN within the image stack.

A normalized fluorescence difference (FD) stack for each slice was then produced by first background subtracting the red and green channel separately for each z plane and then adjusting the red and green pixel saturation to 5% for each z plane within the dLGN convex hull of a slice’s stack. For each pixel the normalized difference between the contralateral marker fluorescence (tdTomato in most cases) and ipsilateral marker fluorescence (usually EGFP) is then calculated to produce the FD (ranging from −1 to1) stack of each brain slice:

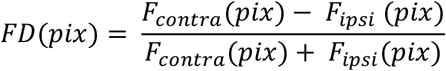

Where F denotes the fluorescence intensity and pix the pixel. In the case of the binarized FD stack a 3D 64 μm standard deviation Gaussian filter was first applied to each channel in the confocal image stack before producing the FD stack. The FD values were then thresholded, such that pixels with FD values over 0 were assigned a value of 1 and those with a value below 0 were assigned a value of −1.

### Alignment of patch-clamp data to ACCF and visuotopic data

The 10 μm voxel resolution ACCF was downloaded from http://data.cortexlab.net/allenCCF/ and contained pixel wise indexes for the dLGN, dLGN shell & dLGN ipsilateral projection zone. To align the position of the cells in our dataset to the ACCF custom-written MATLAB code was used to first manually compare the overall shape of dLGN and that of the ipsilateral projection zones in the ACCF to the confocal image stack of each brain slice. The matching anterior-posterior ACCF section was first determined, and then the position of each cell within that section was identified. During this positioning procedure, the experimenter was blinded to the functional properties and the dendritic morphology of the dLGN neurons. To produce an interpolated/extrapolated 3D map of ODI within dLGN the local weighted average of cell ODIs with 375μm radius around each voxel was calculated used a Gaussian kernel with 75μm standard deviation. To prevent extrapolation artefacts, voxels outside of the sampled anterior-posterior range were not assigned a local average ODI, meaning the anterior and posterior tips of the dLGN were not mapped.

To generate a visuotopic map of dLGN registered to the ACCF, *in vivo* electrophysiological data generously provided to us by the Niell lab^34^ was used. This dataset included 257 units with visuotopic elevation and azimuth receptive field center information as well as their estimated x (medio-lateral) and y (dorso-ventral) position within one of three dLGN schematic outlines from three coronal planes separated in the anterior-posterior axis. Custom-written MATLAB code was used to manually align (via scaling, translation and rotation) these three schematic dLGN outlines to the ACCF. The interpolated and extrapolated 3D map of dLGN azimuth and elevation was produced in the same manner as for the 3D map of average ODI.

To produce a map of average ODI across visuotopy, the mean ODI of all voxels with equivalent elevation and azimuth was calculated. To calculate the average azimuth and elevation gradient within the dLGN, the numerical gradient was first calculated, of either azimuth or elevation, in three dimensions for each voxel within the dLGN. These numerical gradients were then converted to polar coordinates. The gradient angle was defined as the polar component corresponding to the gradient in the coronal plane. The average of all gradient angles across all voxels then gave the average gradient of elevation and azimuth.

### Calculation and comparison of mFD and rFD

For each cell whose location in the confocal image stacks could be re-identified a 600 μm by 600 μm (x by y) column, centered on the soma position, was cut out of the corresponding FD stack. This stack was then down-sampled to a voxel size of 14.6 μm. To calculate mFD, first the interpolated and aligned morphology was used to create a 3D histogram of relative morphology densities. This 3D histogram was then used to calculate the weighted average of FD values based on the voxel wise morphology density for each cell. The mFD calculation was repeated after rotating the dendritic morphology of each cell through 360°, in 10° increments, to obtain mFD at different rotation angles. To estimate the probability of a cell with a given mFD value being contralaterally dominated the percentage of contralateral cells, within +/− 0.2 of the mFD of each cell, was calculated.

To calculate radial mask-based FD (rFD) the same procedure was performed but instead of the cells own morphology, the average 3D dendrite density across all cells was used. The average relative dendritic density histogram, across cells, as a function of Euclidean distance from the soma was calculated and each distance bin was divided by the area that distance bin occupied in 3D space. This histogram was used to calculate the relative dendrite density at each voxel, within the downsampled FD stack. For both rFD and mFD the voxel containing the cell soma was not included in the calculation to avoid contribution of Alexa-594 bleed-through in the red channel.

To obtain the rFD distribution across a whole slice, a grid of sampling points was defined, with 40 μm spacing between points, at the mean z position of cells patched in that slice. The rFD was then calculated for each of these sampling points. This procedure was repeated with the binarized FD stack in order to estimate the predisposition to monocularity based on the size of the contra- and ipsilateral projection zones, to simulate ideal conditions of perfect projection segregation.

To compute the decoding accuracy of mFD the fraction of cells who’s ODI and mFD had the same sign was divided by the total number of cells. To determine if this decoding accuracy was above chance cell ODI was shuffled 10,000x and the decoding accuracy recalculated. The actual decoding accuracy was considered above chance if it was above the upper 5% percentile of the shuffle distribution.

d’ for each of the two masks over 10,000 bootstrapped shuffles (*d*’(*bsh*)) was calculated to compare the discriminability of contra- and ipsilateral dominant cells based on mFD and mFD when morphologies where rotated by 180°:

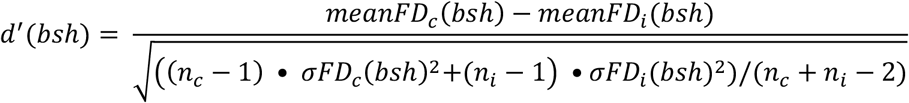

Where c denotes contralateral dominated cells, i ipsilateral dominated cells, bsh the bootstrapped shuffle iteration (1 to 10,000), meanFD the mean FD measure (eg. mFD), σFD the standard deviation of the FD measure and n the number of cells. This distribution was the used to calculate the two-sided p-value of the bootstrapped differences of d’:

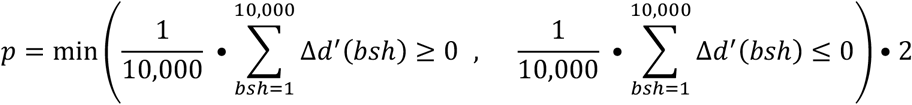

Where Δ*d*’(*bsh*) is the difference in d’ between the two FD measures at each bootstrap iteration.

### Estimation of total contra-, ipsilateral and binocular dLGN neurons across the whole dLGN

The distribution of rFD values across a slice was subsequently used to estimate the overall proportion of dLGN input types. For this, the fraction of 40 μm spaced sampling positions across a slice which fall within a given rFD bin (*sFD_hist_*) was calculated:

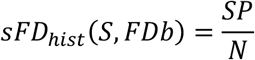

Where S denotes the slice, FDb the rFD bin (six equidistant bins from −1 to 1), SP denotes the number of sampling positions across a slice that fall within the rFD bin and N the total number of sampling positions measured across slice. The mean distribution for each anterior-posterior position bin (*apFD_hist_*; 5 equidistant bins from anterior-posterior position 7.010 mm to 8.210 mm in the ACCF), is defined as:

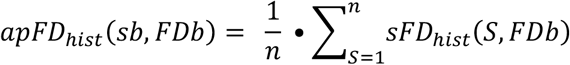

Where sb denotes the anterior-posterior bin and n the number of slices within a given rFD bin. To compensate for varying cross-sectional area at different anterior-posterior slice positions, we calculate the weighted mean distribution across anterior-posterior bins (*W_sa_*(*sb*)) was calculated:

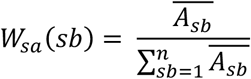

Where 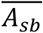 denotes the mean cross-sectional area of all slices within an anterior-posterior bin. Finally, the expected FD distribution across the whole dLGN (*expFD_hist_*) was calculated:

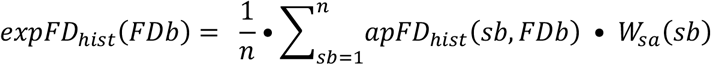

Where n denotes the number of anterior-posterior bins. This process was repeated with the binarized version of the confocal image stack to calculate the fraction of neurons that would get roughly equal proportion of contra- and ipsilateral input, providing the RGC projection zones were perfectly segregated. The ratio between the expected rFD distribution and the observed rFD distribution then gave the correction factor (*CF*):

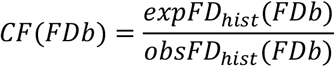

After determining the fraction of neurons with an ODI under −0.333 (ipsiF), over 0.333 (contraF) and between −0.333 and 0.333 (binoF) in each rFD bin the relative contribution of each rFD bin based on the correction factor was adjusted:

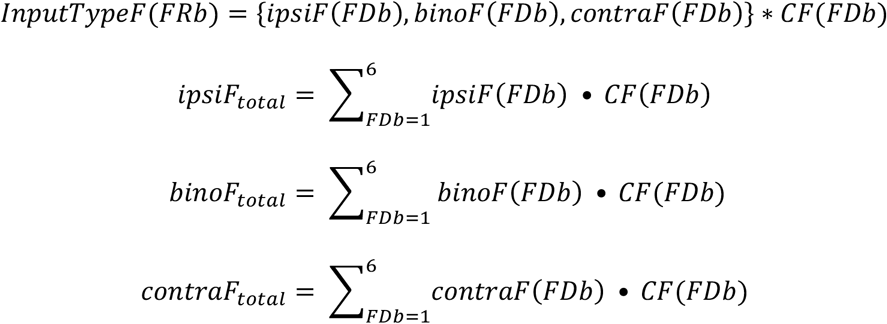

### Calculating background FD gradient

To calculate background FD gradient, a 600 μm by 600 μm (x by y) column around each cell from the down sampled FD stack was again taken, but this time a spherical mask was applied, with radius 150um and centered on the cell soma. Using the FD values in this sphere linear regression was performed, with the displacement in x and y from the cell soma of each pixel as the regressors. The length of the vector formed when x and y are set to the regression coefficients was then taken as the background FD gradient magnitude. The orientation (between −90° and +90°) of the vector was’ first adjusted for the rotational offset between the confocal image stacks and the ACCF schematics and defined as the background FD gradient orientation.

### Quality control and data exclusion criteria

In total, data from 220 dLGN neurons was acquired during our *in vitro* experiments. Two separate semi-automated quality control procedures were performed in which several parameters were checked and judged independently: 1) electrophysiological data quality, 2) confocal imaging and cell re-identification, transduction, dendritic morphology tracing and ACCF alignment quality. During the two quality control procedures the experimenters were blinded to either the electrophysiological data or the imaging data, respectively.

AMPAR responses from 38 cells and NMDAR responses from 67 cells were excluded based on poor quality and an additional 14 cells were excluded when comparing AMPAR and NMDAR due to a series resistance changing by more than 50% between the two recordings. Recordings in voltage-clamp mode of cells patch-clamped with a K-gluconate-based internal solution (26 cells) were only used in Fig. 2h. The dendritic morphology of 34 cells were not used due to poor quality tracing or the total length of dendrites being under 600 μm (an indication of inadequate cell filling with Alexa-594). One cell was additionally excluded from all analyses based on its morphological resemblance to an interneuron^67^. The confocal data of 25 cells was excluded due to poor imaging quality and were not localized in the ACCF. The electrophysiological and fluorescence quantification data of 17 cells were excluded due to insufficient local transduction. Likely due to anisotropic compression of the brain slices during preparation for confocal imaging, the dendrites of some cells reached out of the imaged slice. mFD values of 20 cells with more than 30% of their dendrites protruding out of the slice were not included in analyses based on mFD. A further 18 cells were excluded, that had more than 10% of their dendrites reach out of the dLGN when they were rotated, from analysis using mFD values from rotated morphologies. Individual aspects of the data (e.g. morphology traces, AMPAR responses) were excluded based only on quality control aspects relevant to that data type (e.g. when quantifying morphological features, only criteria relevant to dendritic morphology traces were applied).

For *in vivo* experiments, one animal was excluded due to lack of expression and a second one due to repeated bone growth preventing optical access.

### Statistics

Data are reported as mean ± standard error of the mean (SEM) unless stated otherwise. For circular statistics, the circ_stats^68^ and circStatNP toolboxes (https://github.com/dervinism/circStatNP) were used. Before comparison of data, individual data sets were checked for normality, in the case of linear variables. When normality could not be ruled out, using the Kolmogorov-Smirnov Goodness-of-Fit test, two-tailed two-sample unequal-variance, two-way ANOVA, or paired t-tests were used. When normality was ruled out or could not be assumed, the Wilcoxon rank-sum test, the Mann-Whitney U test or the Kruskal-Wallis test were used. In the case of periodic variables, a von Mises distribution was assumed.

To test for correlations between two linear variables Pearson’s correlation was used when the assumptions of normality could be fulfilled and Spearman’s correlation otherwise. Circular-circular correlation was used to correlate two periodic measures, and circular-linear correlation when comparing a periodic and a linear measure. Here, Pearson’s correlation was also used when the assumptions of normality (or a von Mises distribution in the case of periodic variables) could be assumed and Spearman’s when they could not. If the maximum possible range of a periodic variable was 180° rather than 360°, we multiplied the angle by 2 before converting to radians and performing statistical tests for correlations in order meet the assumptions of a von Mises distribution.

Asterisks indicate significance thresholds as follows: p < 0.05 (*), p < 0.01 (**), p < 0.001 (***), unless otherwise stated. p-value thresholds were adjusted using the Bonferroni method for multiple comparisons correction where necessary.

## Data availability statement

The datasets generated and analyzed during the current study are available from the corresponding author upon request.

## Code availability statement

Custom code developed for analyzing the data during the current study is available upon request.

## Acknowledgements

We are grateful to Volker Staiger, Meike Hack and Ursula Weber for cell tracing as well as Volker Staiger for excellent technical support, to Michael Myoga for helping to build one *in vitro* setup, to Robert Kasper for assisting with confocal imaging, Sandra Reinert for useful discussions, Pieter Goltstein for software. Additionally, we are grateful to Botond Roska, Josephine Juettner, and Juliane Jäpel for help with intravitreal injections. This study was supported by the Max Planck Society and the Deutsche Forschungsgemeinschaft (CRC 870; T.R., V.S., M.H., and T.B.).

## Author Contribution

S.W., J.B., M.F. and T.R. conceived the project; M.F. and S.W performed electrophysiological experiments. D.L. performed *in vivo* experiments. J.B. and S.W. wrote *in vitro* analysis tools and analyzed the corresponding data. D.L. and T.R. analyzed the *in vivo* data. J.B. performed intravitreal eye injections. M.F. performed confocal imaging and cell re-identification. D.L. developed the viral construct for *in vivo* imaging. V.S. assisted with dual-channel input mapping. T.R., J.B., S.W., M.F., D.L., M.H., V.S, and T.B. wrote the manuscript. All Authors discussed the data. T.B. provided the research environment.

## Affiliations

Max Planck Institute of Neurobiology, Martinsried, Germany: Joel Bauer, Simon Weiler, Martin Fernholz, David Laubender, Volker Scheuss, Mark Hübener, Tobias Bonhoeffer, Tobias Rose

## Corresponding author

Correspondence to Tobias Rose

