## Supplementary Figures for "Selective connectivity limits functional binocularity in the retinogeniculate pathway of the mouse"

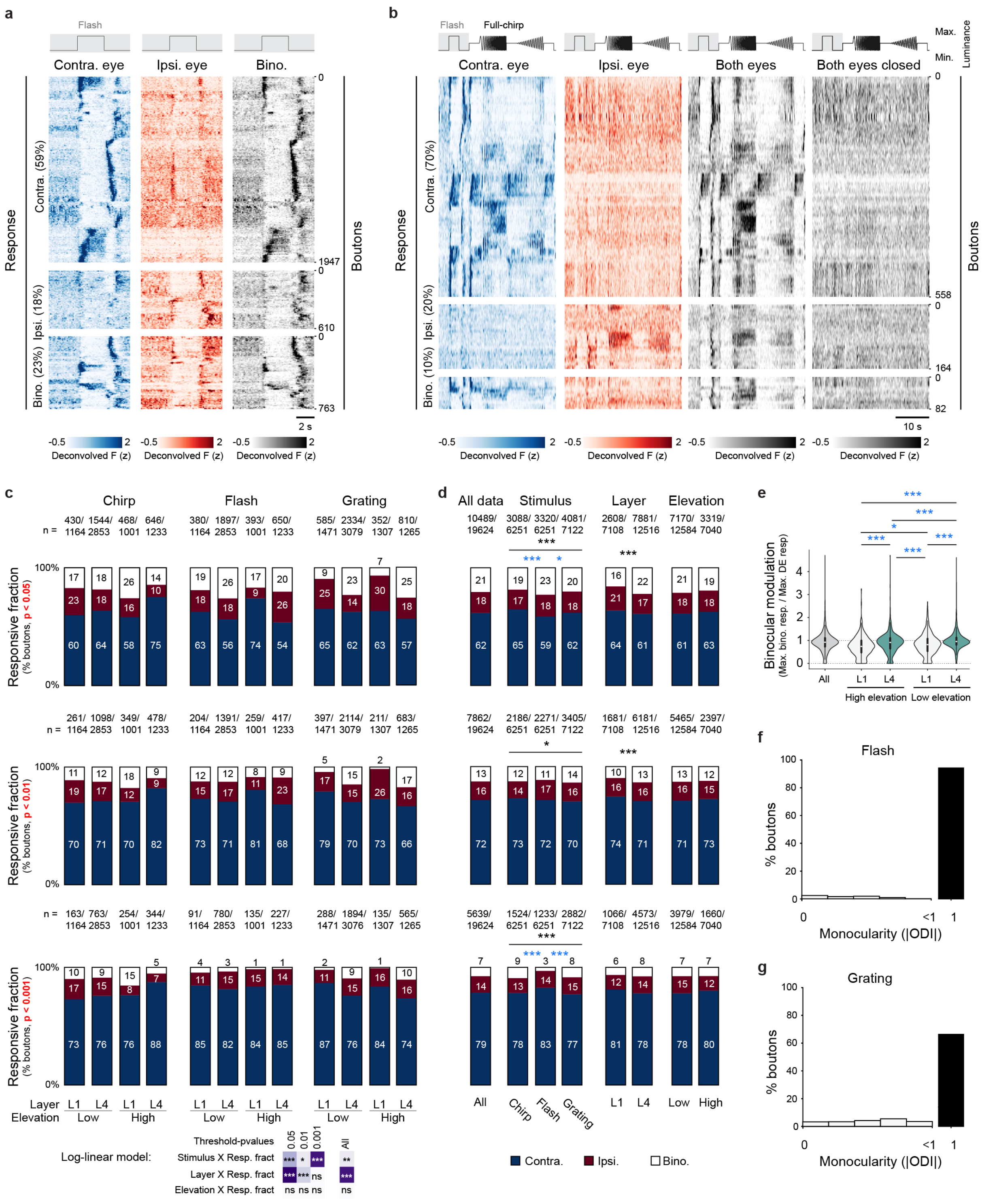


### Supplementary Fig. 1: Eye-specific thalamocortical neuron binocularity in relation to stimulus type, visuotopy, and cortical projection layer

**a**, Activity of 3320 boutons (N = 4 mice) across layers and elevations in response to contralateral, ipsilateral, and both-eye full-field luminance stimulation, grouped by eye-specific responsiveness and sorted by similarity of the joint stimulus-aligned data (boutons with highly correlated responses lie next to each other, see methods). Data as in Fig. 1c but selected, sorted, and displayed based on responses to the initial full-field flash part of the full-length luminance stimulus (including both significant positive and negative responses). **b**, as in a (n = 804 boutons, N = 3 mice) but in response to the full-length luminance stimulus and including stimulation with both eyes closed to assess light leak through the eye shutters. Note lack of responses under this condition. **c**, Eye-specific response fractions across all conditions with three different threshold criteria (p-threshold of $p < 0.05$, top, $p < 0.01$, middle, $p<0.001$, bottom; including significant positive and negative responses; number of responsive boutons over total number of boutons indicated on top of bars, percentages indicated in stacked bar subgroups); Bottom panel, left: p-value table of log-linear model single p-threshold of 0.05: 4 factor single pairwise interaction vs. non-interaction main model (Residual deviance = 781, df = 29): Stimulus X Response fraction (Residual deviance = 753, df = 25), $\chi^2(2)$ $p < 0.001$; Layer X Response fraction (Residual deviance = 724, df = 27), $\chi^2(2)$ $p < 0.001$; Elevation X Response fraction (Residual deviance = 776, df = 27), $\chi^2(2)$ $p = 0.09$; log likelihood-ratio model comparison using $\chi^2$ test; log-linear model single p-threshold of 0.01: 4 factor single pairwise interaction vs. non-interaction main model (Residual deviance = 669, df = 29): Stimulus X Response fraction (Residual deviance = 656, df = 25), $\chi^2(2)$ $p = 0.014$; Layer X Response fraction (Residual deviance = 650, df = 27), $\chi^2(2)$ $p < 0.001$; Elevation X Response fraction (Residual deviance = 668, df = 27), $\chi^2(2)$ $p = 0.43$; log-linear model single p-threshold of 0.001: 4 factor single pairwise interaction vs. non-interaction main model (Residual deviance = 601, df = 29): Stimulus X Response fraction (Residual deviance = 541, df = 25), $\chi^2(2)$ $p < 0.001$; Layer X Response fraction (Residual deviance = 595, df = 27), $\chi^2(2)$ $p = 0.06$; Elevation X Response fraction (Residual deviance = 596, df = 27), $\chi^2(2)$ $p = 0.09$; Bottom panel, right: p-value table of log-linear model across p-thresholds: 5-factor selected dual pairwise interaction (always including Response fraction X p-threshold) vs. single-interaction Response fraction X p-threshold main model (Residual deviance = 2404, df = 95): Stimulus X Response fraction (Residual deviance = 2386, df = 91), $\chi^2(2)$ $p = 0.0014$; Layer X Response fraction (Residual deviance = 2353, df = 93), $\chi^2(2)$ $p < 0.001$; Elevation X Response fraction (Residual deviance = 2399, df = 93), $\chi^2(2)$ $p = 0.09$; log likelihood-ratio model comparison using $\chi^2$ test; **d,** as in c but pooled over the independent variables of the experiment (All data across p-thresholds, 3x3 $\chi^2(4) = 668$, $p < 0.001$; mean absolute deviation of binocular fraction (MADbino) = 6%; post-hoc test p-threshold = 0.001 vs. p-threshold of 0.05 $p < 0.001$; p-threshold of 0.01 vs. p-threshold of 0.05 $p < 0.001$; p-threshold of 0.01 vs. p-threshold of 0.001 $p < 0.001$; Stimulus at p-threshold of 0.05, 3x3 $\chi^2(4) = 28$, $p < 0.001$; MADbino = 2%; post-hoc test Chirp vs. Grating $p = 0.13$; Chirp vs. Flash $p < 0.001$; Flash vs. Grating $p = 0.021$; Layer at p-threshold of 0.05, 2x3 $\chi^2(2) = 56$, $p < 0.001$; MADbino = 1.4%; Elevation at p-threshold of 0.05, 2x3 $\chi^2(2) = 5$, $p = 0.09$; MADbino = 0.8%; Stimulus at p-threshold of 0.01, 3x3 $\chi^2(4) = 12$, $p = 0.016$; MADbino = 1%; post-hoc test Chirp vs. Grating $p = 0.26$; Chirp vs. Flash $p = 0.09$; Flash vs. Grating $p = 0.11$; Layer at p-threshold of 0.01, 2x3 $\chi^2(2) = 18$, $p < 0.001$; MADbino = 1.9%; Elevation at p-threshold of 0.01, 2x3 $\chi^2(2) = 1.7$, $p = 0.43$; MADbino = 0.7%; Stimulus at p-threshold of 0.001, 3x3 $\chi^2(4) = 50$, $p < 0.001$; MADbino = 2.2%; post-hoc test Chirp vs. Grating $p = 0.49$; Chirp vs. Flash $p < 0.001$; Flash vs. Grating $p < 0.001$; Layer at p-threshold of 0.001, 2x3 $\chi^2(2) = 5$, $p = 0.064$; MADbino = 0.8%; Elevation at p-threshold of 0.001, 2x3 $\chi^2(2) = 5$, $p = 0.09$; MADbino = 1.1%; all post-hoc tests Bonferroni corrected). **e**, Binocular modulation (Maximum binocular response / Maximum DE response) of visual responses to both-eye vs. single-eye moving grating stimulation across cortical thalamorecipient layers and visuotopic elevations (two-way ANOVA: main effects: Elevation $F(1,47)$ $p < 0.001$, Layer $F(1,94)$ $p < 0.001$, interaction: Elevation X Layer $p = 0.45$; post-hoc tests: High/L1 vs. High/L4 $p < 0.001$; Low/L1 vs. Low/L4 $p < 0.001$; High/L1 vs. Low/L1 $p = 0.018$; High/L4 vs. Low/L4 $p < 0.001$; High/L1 vs. Low/L4 $p < 0.001$; High/L4 vs. Low/L1 $p < 0.001$; Mann-Whitney U test, Bonferroni corrected). **f**,**g,** Monocularity (|ODI|) of data in response to full-field flash (f, mean |ODI| = 0.95 ± 0.18, positive responses only) and moving grating stimulation (g, mean |ODI| = 0.9 ± 0.23; $w = 3e^{-14}$; Mann-Whitney U tests; positive responses only).

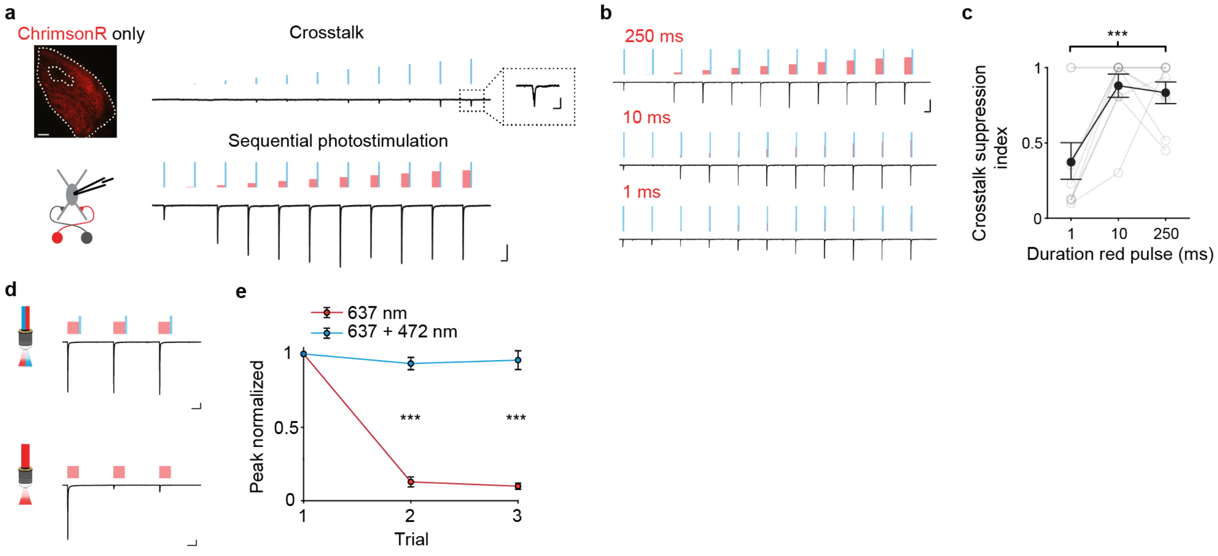


**Supplementary Fig. 2: Sequential photostimulation is necessary for spectral crosstalk suppression and rundown prevention of ChrimsonR-evoked currents**

**a,** Maximum intensity projection of expression pattern of ChrimsonR-tdT within dLGN. Top: PSCs evoked by stimulation at 473 nm using 11 steps with increasing irradiance. Inset, zoom-in of evoked PSC at last step. Bottom, Evoked PSCs of the same dLGN neuron using sequential stimulation at 637 nm (250 ms) and 473 nm (50 ms, ISI: 10 s) **b,** PSCs evoked by 250, 10 or 1 ms stimulation at 637 nm using 11 steps with increasing irradiance paired with constant 50 ms stimulation at 473 nm. **c,** Comparison of average crosstalk suppression index for 1, 10 and 250 ms stimulation at 637 nm (two-way ANOVA, main effect: $p < 0.01$; post-hoc test: 1 ms vs 10 ms: $p < 0.01$, 1 ms vs 250 ms: $p < 0.01$, 10 ms vs. 250 ms: $p = 0.94$). **d,** PSCs evoked by repetitive stimulation at 673 nm with (top) and without paired stimulation at 473 nm (bottom). **e,** Average peak normalized responses of ChrimsonR evoked by 250 ms stimulation at 637 nm without and with paired stimulation at 473 nm for individual repetitions (two-sample student t-test, trial 2 with blue vs. trial 2 without blue: $p < 0.001$, trial 3 with blue vs. trial 3 without blue: $p < 0.001$). Error bars indicate mean ± SEM. Image scale bar: 100 µm. Electrophysiology trace scale bars: 200 ms, 250 pA (zoom in window in **a**: 50 ms, 75 pA).

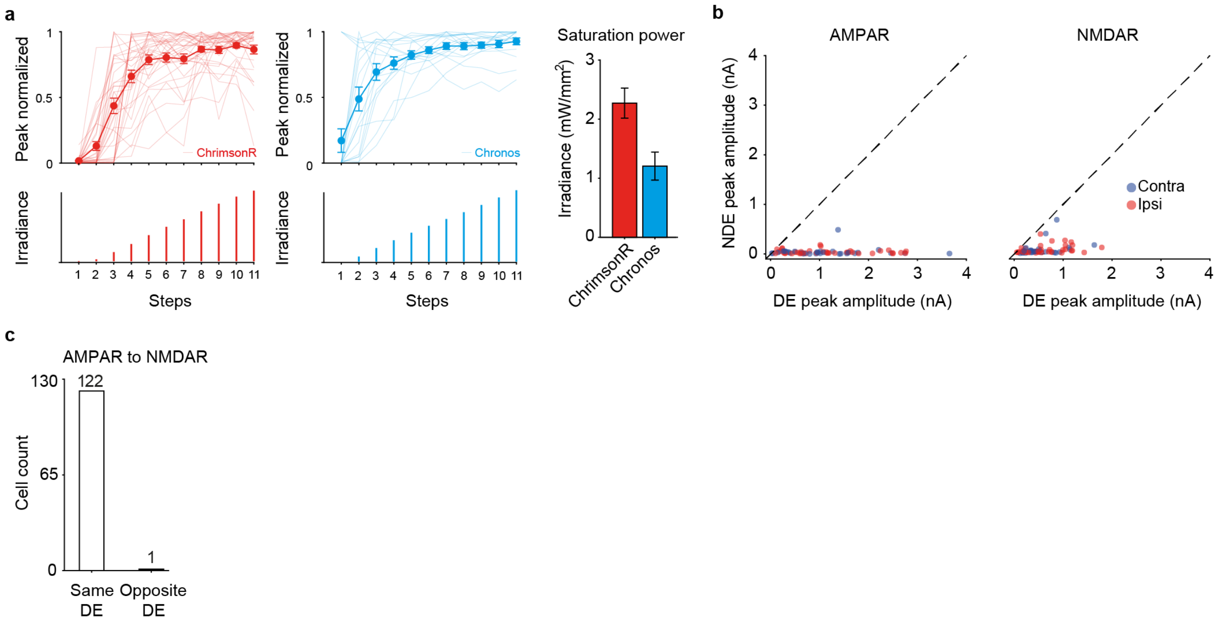


### Supplementary Fig. 3: Assessing dominant and non-dominant eye responses using dual-channel CRACM

**a,** Left**,** average peak-normalized responses of ChrimsonR evoked by stimulation at 637 nm using 11 steps of increasing irradiance. Thin lines indicate individual cells, thick line mean ± SEM. Right, average peak-normalized responses of Chronos evoked by stimulation at 473 nm using 11 steps of increasing irradiance. Thin lines indicate individual cells, thick line mean. Far right, average irradiance used for ChrimsonR and Chronos to obtain current saturation. **b,** Evoked peak currents of the DE plotted against the evoked peak currents of the NDE for AMPAR- (left, n = 75) and NMDAR- mediated (right, n = 65) responses for binocular cells. Color indicates whether the contra- or ipsilateral eye is dominant. **c,** Comparison of dominant eye preference for AMPAR- and NMDAR-mediated responses (n = 123 cells). Error bars indicate mean ± SEM.

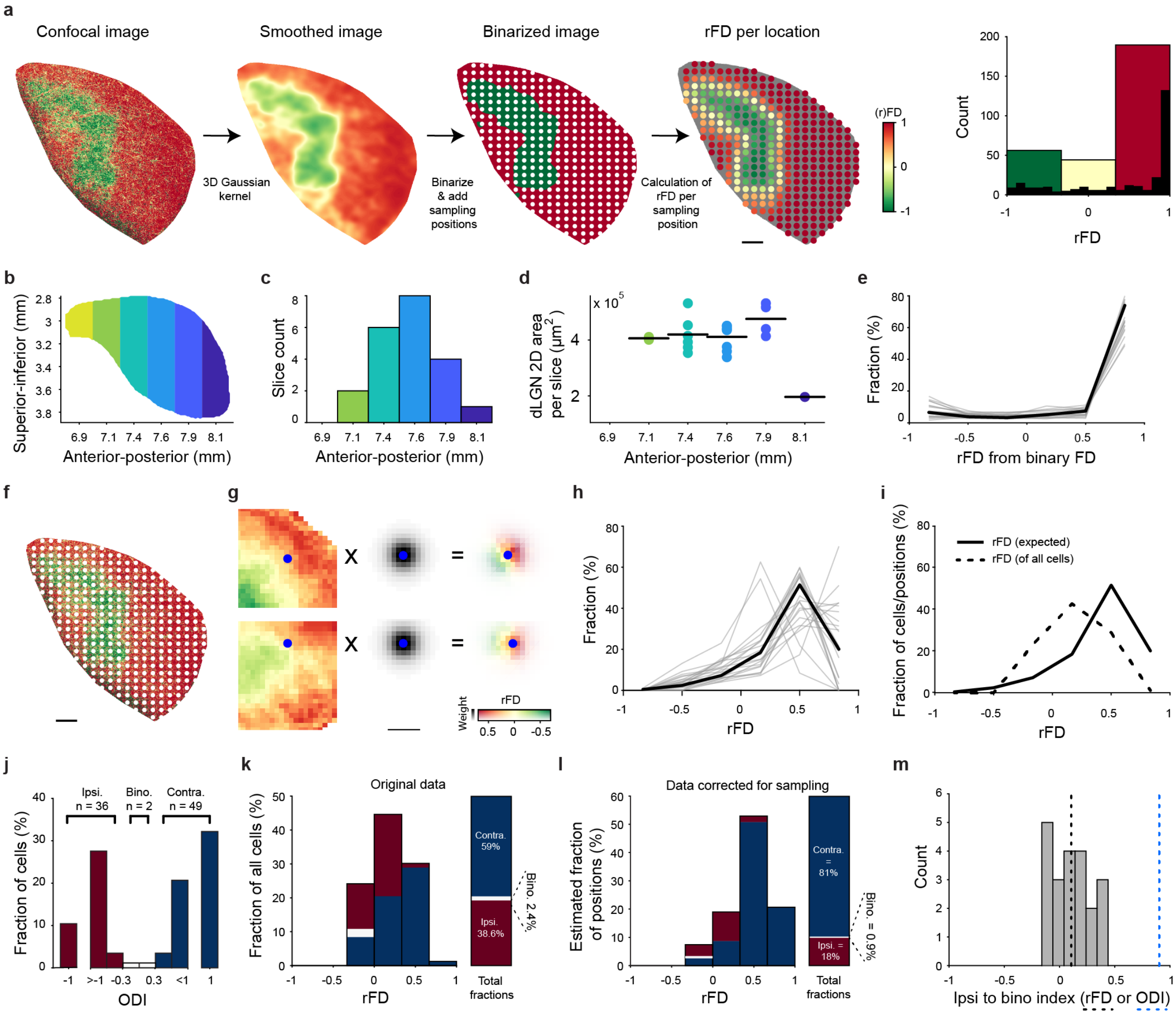


### Supplementary Fig. 4: Calculation of the total fraction of binocular neurons in dLGN using rFD

**a,** Processing of example slice for obtaining estimated rFD distributions under conditions of perfect axonal segregation. From left to right: fluorescence difference map of single plane from confocal stack, smoothing using 3d Gaussian kernel, image binarization, indication of sampling positions in white, rFD values for each sampling position (dLGN in gray) and distribution of mFD separated into three equidistant bins (ipsi: mFD -1 to -0.333, bino: mFD -0.333 to 0.333, contra: mFD 0.333 to 1). Black bars: 30 bin histogram. **b,** ACCF based dLGN schematic with indicated anterior-posterior bins (bin color same for **c** and **d**). **c,** Histogram of slice number per anterior-posterior bin (n = 21 slices). **d,** Surface area of dLGN in each slice and mean for each anterior-posterior bin. **e,** Line histogram of rFD samples across each binarized slice (gray) and weighted mean across all slices (black). Weighted mean takes slice frequency and surface area of each anterior-posterior bin into account. **f,** Fluorescence difference of example slice with sampling positions indicated in white. **g,** Process of calculating rFD for example cell location. Left to right: down-sampled fluorescence ratio stack, mean radial morphology mask, and masked fluorescence ratio pixel map based on mean radial morphology mask. Top row displays top view of coronal brain slice; bottom row displays side view of coronal brain slice. Cell soma location indicated by blue circle. **h,** Same as for **e**, but for non-binarized fluorescence slices. **i,** Distribution of rFD values of cells patched across all slices (dashed line) and expected rFD distribution given even sampling across and between slices (solid line, same as in **h**; n = 87 cells, 21 slices). **j,** ODI distribution, indicating the three equidistant ODI bins (red: ODI >= -1 & < 0.333; white: ODI >= -0.333 & <= 0.333; blue: ODI > 0.333 & <= -1). **k,** Distribution of ODI fractions (indicated in **j**) across rFD bins and as a fraction of total cell count (n = 87 cells). **l,** Same as **i**, but relative weighting of each rFD bin is adjusted to match the expected rFD distribution and total corrected fraction. Given the total number of excitatory cells in dLGN (~20000; [46]), we therefore estimate that there are ~180 neurons that have a DE to NDE input ratio of over 0.5 (|ODI| < 0.333) in the mouse dLGN. **m,** Summary histogram showing distribution of normalized difference between ipsilateral and binocular sampling positions of each slice (gray bars), the weighted average (dashed black line) and the normalized difference of functionally ipsilateral and binocular neurons (dashed blue line). All scale bars: 100 µm.

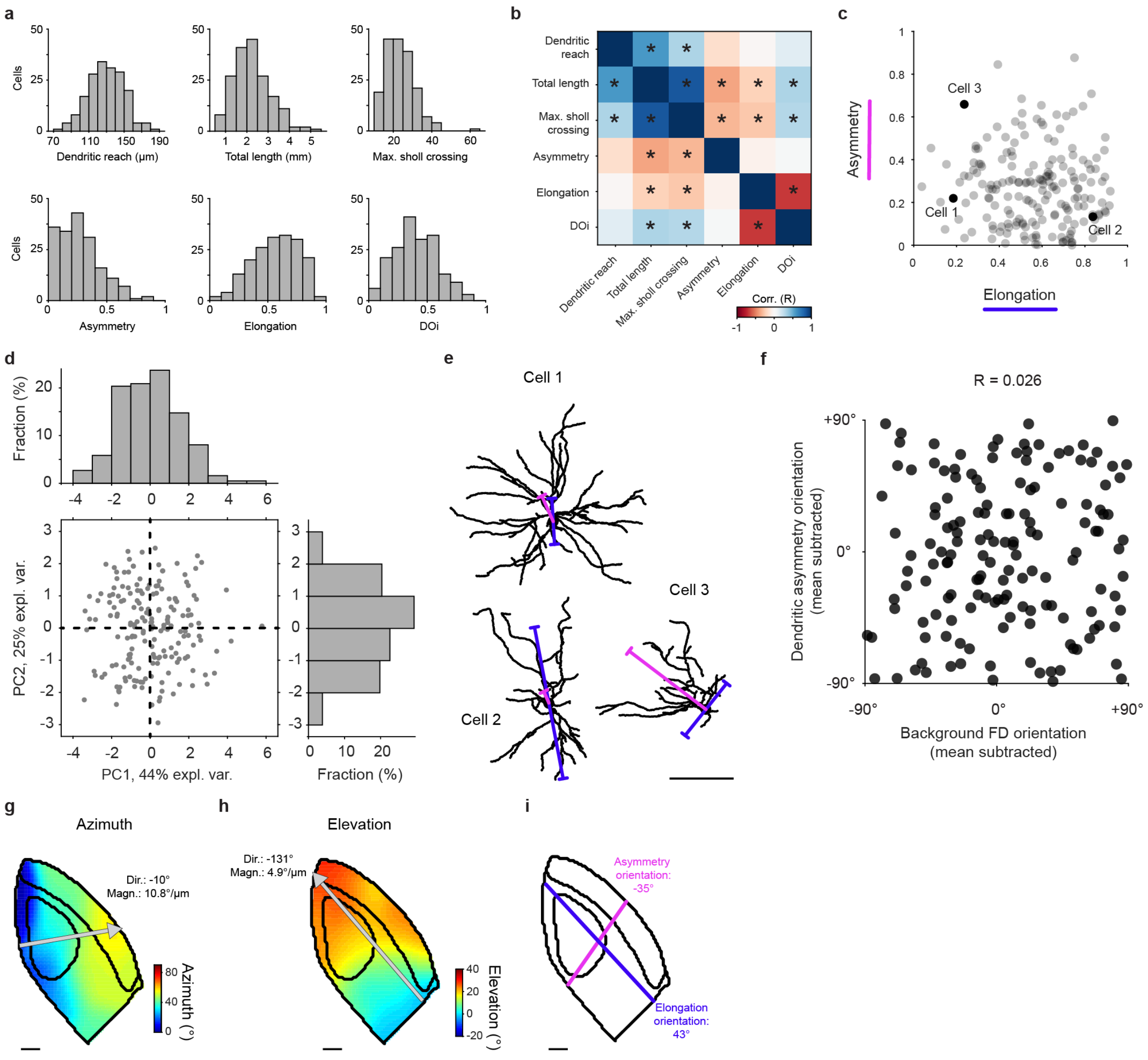


### Supplementary Fig. 5: dLGN dendritic morphology primarily varies in reach, elongation, and asymmetry

**a,** Distributions of six morphological features: dendritic reach, total dendritic length, Max. number of crossings (Sholl analysis), asymmetry magnitude, elongation magnitude, and DOi (n = 185 cells). **b,** Spearman's correlation coefficient matrix for the six morphological measures. * indicates $p < 0.05$ after Bonferroni correction. **c,** Magnitude of asymmetry versus elongation of all cells (n = 185 cells). Example cells in **e** are indicated in black. **d,** Distributions of all dLGN neurons along the first two principal components. Histograms of both components displayed on top and left of the scatter plot. Neither components pass the dip test for multimodality (n = 185 cells, first principal component Hartigan's dip index = 0.023, p = 0.76; second principal component Hartigan's dip index = 0.019, p = 0.96); the data is therefore not suitable for clustering analysis. **e,** Dendritic morphology of three example dLGN neurons. Lines indicate angle and magnitude of the morphological parameters asymmetry (magenta) and elongation (blue). Lines representing elongation are bidirectional because elongation angle is only defined in terms of orientation and has no direction. The length of the line indicates the product of the magnitude of elongation or asymmetry, and the dendritic reach from soma. **f,** Scatter plot of mean subtracted background FD gradient orientation vs mean subtracted asymmetry orientation (n = 152 cells, circular-circular Pearson's correlation R = 0.026, p = 0.74). **g,** dLGN single slice schematic (based on ACCF) with interpolated visuotopic azimuth map and average gradient of azimuth change (calculated in 3D, gray arrow indicating coronal plane component; dir = -10°, Δazimuth per µm = 10.8 °/µm). **h,** Same as in **g** but for elevation (dir = 131°, Δelevation per µm = 4.9 °/µm). **i,** Mean dendritic morphology asymmetry and elongation angles of dLGN neurons (asymmetry: ori = -35°; elevation: ori = 43°). Interestingly, we found that the mean elongation angle of dLGN neuron dendrites closely aligned with the average visuotopic elevation gradient in **h**. Given that the representation of elevation, relative to azimuth, is stretched in the dLGN this suggests that dendritic elongation may act as a compensatory mechanism for this anisotropy in visual mapping. All scale bars: 100 µm.

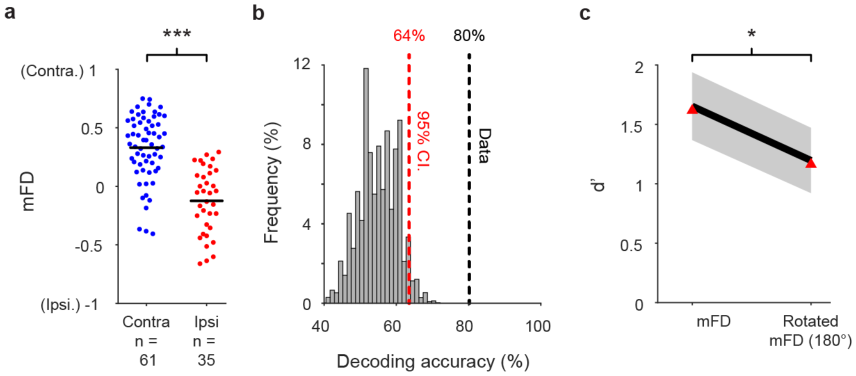


**Supplementary Fig. 6: Decoding eye preference from local RGC innervation**

**a,** Distribution of mFD values for contra- and ipsilaterally dominated cells (Contralateral = 61 cells, ipsilateral = 35 cells, two-tailed two-sample unequal variance t-test $p < 0.001$, $t = 7.7$). **b,** Decoding accuracy of eye preference from mFD (black dashed line), histogram of decoding accuracies of shuffled eye preference with 95% confidence interval indicated (red dashed line). **c,** d' comparison of mFD, at 0° and 180° rotation from original orientation. Red triangles: actual d' based on data, gray: standard deviation of the bootstrapped d' distributions, black: mean of bootstrapped d' distributions (n = 82 cells, mFD vs mFD at 180° rotation: two-tailed paired bootstrap p-value of the difference between d' values: $p = 0.0142$).
